# Dynamic signatures of lipid droplets as new markers to quantify cellular metabolic changes

**DOI:** 10.1101/2020.07.28.225235

**Authors:** Chi Zhang, Stephen A. Boppart

## Abstract

The metabolic properties of live cells are very susceptible to intra- or extra-cellular perturbations, making their measurements challenging tasks. We show that the dynamics of lipid droplets (LDs) carry information to measure the lipid metabolism of live cells. Coherent anti-Stokes Raman scattering microscopy was used to statistically quantify LD dynamics in living cells in a label-free manner. We introduce dynamic signatures of cells derived from the LD displacement, speed, travel length, and directionality, which allows for the detection of cellular changes induced by stimuli such as fluorescent labeling, temperature change, starvation, and chemical treatment. Histogram fittings of the dynamic signatures using lognormal distribution functions provide quantification of changes in cellular metabolic states. The LD dynamics also enable separation of subpopulations of LDs correlated with different functions. We demonstrate that LD dynamics are new markers to quantify the metabolic changes in live cells.

## Introduction

Metabolic homeostasis is the key to maintain proper functions of living cells, while altered metabolism is usually associated with disease transitions. Among different metabolites, lipids are crucial for cellular energy storage, membrane construction, signaling, and hormone production. Lipid droplets (LDs), a key organelle that is responsible for the storage and regulation of neutral lipids in cells, has stepped into the spotlight of lipid research in the past two decades due to its direct association with cancer transition and progression (Bozza and Viola, 2010; Petan et al., 2018; Tirinato et al., 2017). Scientists realized that LDs are not simple and inert lipid deposits, but are highly dynamic and functional structures which are major lipid regulators in cells (Baenke et al., 2013; Walther and Farese Jr, 2012). Recent studies found that LD biogenesis is directly linked to cellular stress factors such as reactive oxygen species (Bailey et al., 2015; Welte, 2015) and hypoxia (Biron-Shental et al., 2007; Koizume and Miyagi, 2016), and offers protective effects for cells in stress environments (Bosma et al., 2014; Herms et al., 2015; Jarc et al., 2018). This highlights the critical roles of LDs in regulating cell metabolism to maintain cellular homeostasis under various external stimuli. Besides, LDs were found to accumulate in various cancer cells (Mitra et al., 2017; Morjani et al., 2001), suggesting their indispensable association with metabolic reprogramming in cancer. LDs were reported to protect cancer cells from the treatment of chemotherapeutic drugs (Cotte et al., 2018) and accumulate cholesteryl ester to promote tumor growth (Yue et al., 2014). Elucidating LD metabolism has led to discoveries of new targets for cancer treatment (Liu et al., 2017; Yue et al., 2014) and cancer stem cell suppression (Li et al., 2017).

Despite numerous efforts that have investigated LD content, size, and amount associated with various stimuli and disease transitions, our understanding of LD dynamics remains primitive. Especially, the dynamic information of LDs, which is an important signature to probe cell metabolism, is largely underexplored due to the lack of appropriate techniques. To identify and distinguish LDs from other organelles, chemical labeling is typically a requisite. Different from other parameters, the dynamic information can only be probed in living samples, ruling out many widely used labeling and imaging techniques such as immune-fluorescence imaging and electron microscopy. Lipophilic dyes such as BODIPY can label LDs in live cells. However, LD dynamics are very sensitive and susceptible to cellular functions, which are perturbed by the introduction of exogenous fluorescent probes. A label-free microscope with lipid selectivity is the desired tool to measure LD dynamics. Mid-infrared and Raman spectroscopic imaging, which are based on vibrational absorption and scattering, have been used to image lipid contents in cells (Pleitez et al., 2020; Zhang et al., 2016). Mid-IR imaging cannot compete with Raman imaging in terms of penetration depth and spatial resolution (Cheng and Xie, 2015), while spontaneous Raman imaging typically has much slower imaging speed on the order of minutes to hours, which is too slow to capture dynamic signatures of LDs in living cells (Palonpon et al., 2013). Coherent Raman scattering microscopy has offered new ways to probe LDs in living cells in a label-free manner with sub-micron resolution and fast imaging speed (Camp Jr et al., 2014; Cheng et al., 2002; Di Napoli et al., 2014; Evans et al., 2005; Freudiger et al., 2008; Fu, 2017; Fu et al., 2012; He et al., 2017; Liao et al., 2015; Lu et al., 2015; Min et al., 2011; Ozeki et al., 2012; Potma et al., 2002; Slipchenko et al., 2009; Wang et al., 2013; Zhang et al., 2013; Zumbusch et al., 1999). With this approach, LD dynamics can be visualized in real-time (Bradley et al., 2016; Nan et al., 2006; Rinia et al., 2008; Zhang et al., 2017).

Although some initial studies have been performed using coherent anti-Stokes Raman scattering (CARS) (Nan et al., 2006) or stimulated Raman scattering (SRS) to measure LD dynamics (Zhang et al., 2017), the comparison of LD trafficking under different conditions is largely qualitative and preliminary. In a previous study, results show a correlation between LD displacement and lipogenesis (Zhang et al., 2017). However, quantitative ways to measure and compare LD dynamics under various conditions are still lacking, preventing the systematic understanding of LD dynamics associated with different environmental stimuli and drug treatment. In this study, we introduce dynamic signatures of living cells by quantification of LD displacement, speed, travel length, and directionality using CARS microscopy. We also used lognormal distributions to model the statistics of LD dynamics, which allows for a quantitative comparison of changes in LD dynamics related to stimuli and drug treatment. We systematically studied LD dynamic responses to fluorescent labeling, hypothermia exposure, apoptosis, starvation, adenosine monophosphate activated protein kinase (AMPK) activation, and drug treatments targeting different lipid metabolic enzymes. We found reduced directionality of LD trafficking during hypothermia exposure and apoptosis, and changes in the ratio of synthesis- and degradation-related LDs during starvation, AMPK activation, and fatty-acid-related enzyme inhibition. Collectively, LD dynamics captured and quantified by CARS imaging offer the potential for a better understanding of biological processes. The LD dynamics might also function as new biomarkers for cellular responses to disease transitions and drug treatments. Our approach allows biologists to understand real-time cell metabolism from a novel angle, and would have important applications in drug development and cancer research.

## Results

### Quantification of LD dynamics

CARS microscopy was used to image LDs in living cells. We tuned the pump and Stokes beams to excite the lipid CH_2_ vibration centered at 2884 cm^-1^. The lab-built CARS microscope used in this study was pumped by a dual-output pulsed laser source and constructed based on an upright microscope frame (Figure S1). The details of our CARS microscope can be found in Materials and Methods. Briefly, the pump and Stokes beams at the sample were ∼ 14 mW, 1 ps, and ∼ 10 mW, 2 ps, respectively. CARS signals were collected in the transmission direction (Figure 1A). Two-photon excitation fluorescence (TPEF) signals from the sample can be simultaneously acquired in the epi direction if needed. The pixel dwell time used for imaging was 10 µs, and the imaging speed for images with 400×400 pixels was 2.2 s/frame. A 100-frame image stack was acquired for each measurement and multiple regions were imaged for each condition (Figure 1B). A Particle Tracker ImageJ Plugin (Sbalzarini and Koumoutsakos, 2005) was used to trace the trajectories of individual LDs that were detected in more than 20 frames. A sample LD trajectory is displayed in Figure 1C and Video S1. A MATLAB-based lab-written program was used to quantitatively calculate all the essential parameters of the trajectories used for statistical analysis.

**Figure 1.**
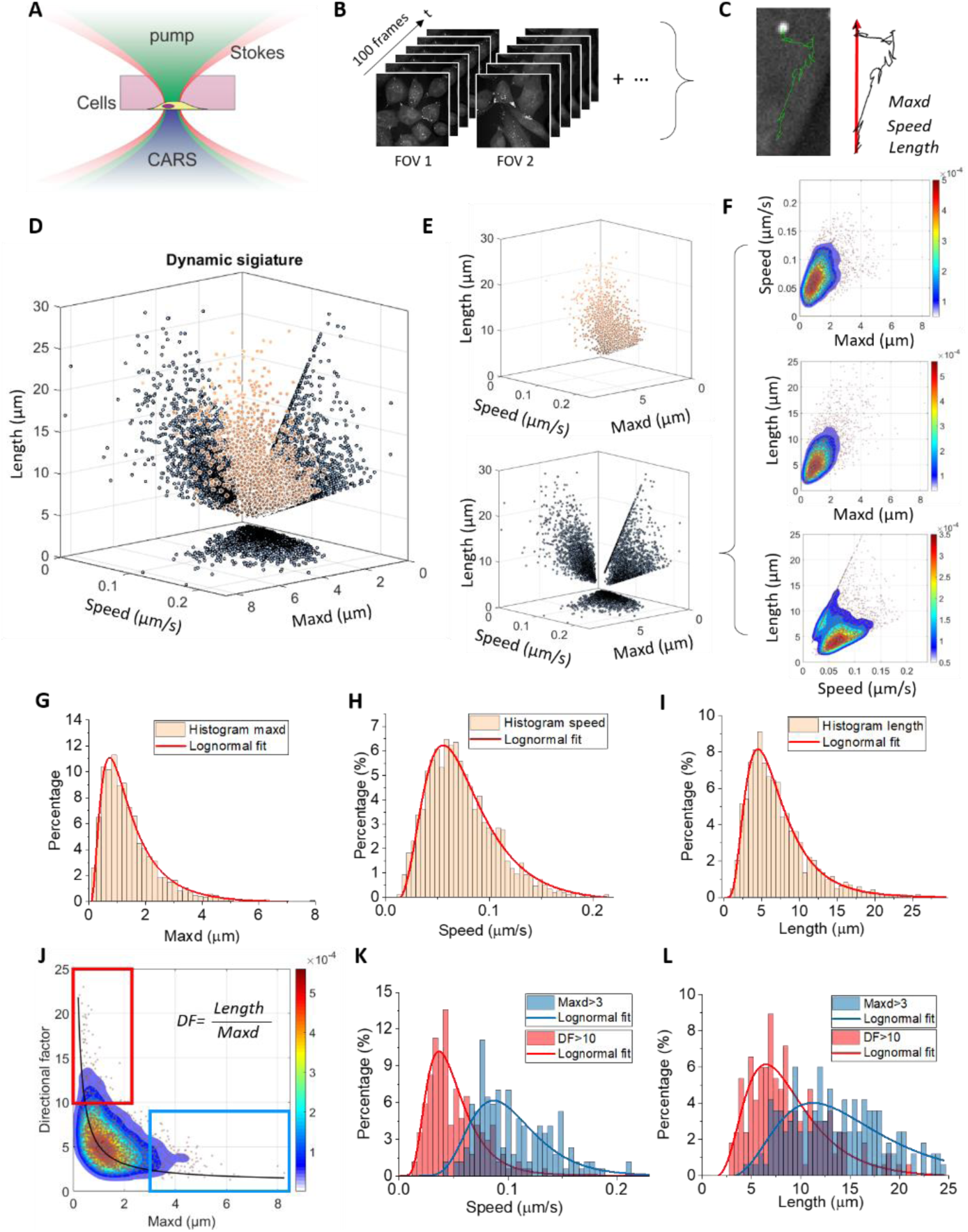
Coherent Raman scattering microscopy for quantitative LD dynamics study. (A) Schematic of CARS signal generation by living cells. (B) Time-lapse imaging of live cells using CARS. (C) An LD trajectory and the parameters derived directly from the trajectory, including the maximum displacement (*maxd*), the average speed, and the total trajectory length. (D) Dynamic signature of MIAPaCa-2 cells plotted in the *maxd, speed*, and *length* 3D space. (E) The 3D plot of the MIAPaCa-2 dynamic signature without (upper panel) projections, and the projections onto the three 2D planes (bottom panel). (F) Density contour plot of the dynamic signature projected onto the *speed-maxd, length-maxd*, and *length-speed* planes. The dots are individual LD trajectories. (G)-(I) One-dimensional histograms of the MIAPaCa-2 dynamic signature projected onto the *maxd, speed*, and *length* axes, respectively, and the lognormal fitting of the histograms. (J) Density contour plot of the dynamic signature projected onto the *DF-maxd* plane. The dots are individual LD trajectories. The black curve is the fitting curve using the reciprocal function. (K) and (L) Histograms and lognormal fitting results of the gated trajectories in panel (J). Red plots are trajectories with *DF* > 10, while blue plots are trajectories with *maxd* > 3.

Three parameters that can be further used to quantify the LD dynamics were directly derived from the trajectories, including the *maxd*, the *speed*, and the *length*. The *maxd* value is defined as the maximum displacement of the LD movement during the time of image acquisition. Different from a previous study which only measured the maximum displacement from the starting point (Zhang et al., 2017), we take the maximum value of the maximum displacement starting from all frames as the *maxd* value (Figure S2). This is a more valid way to quantify the maximum relocation distance of LDs within the time of measurement. The value *length* is the total trajectory length, and the value *speed* is defined as *length* divided by the total time of the trajectory.

The LD trajectories can be plotted in the *maxd-speed-length* 3D space as a signature to characterize LD dynamics of living cells. Figures 1D,E plot the 3D LD dynamic signature of MIAPaCa-2 pancreatic cancer cells (brown dots) at 37 °C in Dulbecco’s Modified Eagle Medium (DMEM) ((+)glucose) + 10% fetal bovine serum (FBS) + 1% penicillin-streptomycin antibiotics. The projection of this 3D dynamic signature onto the three 2D planes (black dots), including *speed-maxd, maxd-length*, and *length-speed*, can be used to better display and compare the changes in LD dynamics. Density contour plots of the dynamic signature in the three planes are shown in Figure 1F, which highlight the population density of the trajectories. Further, the projections of the dynamic signature on the *maxd, speed*, and *length* axes (Figures 1G-I) reveal histograms of the trajectories and can be compared quantitatively through the mathematical fitting. We found that the lognormal function is the best fitting function to quantify these histograms (Figures 1G-I).

In addition to the three parameters defined above, we also derived a ‘*directional factor* (*DF*)’ to quantify the directionality of the trajectories. The *DF* is defined as the total trajectory *length* divided by the *maxd*. A more directional LD trajectory will have a smaller *DF* value. We plotted the MIAPaCa-2 LD dynamics in the *DF-maxd* plane and found a reciprocal relationship between the two parameters (Figure 1J). This indicates that the LDs having longer *maxd* tend to be more directional in their movement, while the LDs having a shorter *maxd* were less directional. This can be explained by the two types of LD dynamics in living cells: LDs traveling in the cytosol are transported along microtubules and are more directional with a longer displacement; while LDs bound to the endoplasmic reticulum (ER) have less freedom to move and are much less directional (Zhang et al., 2017).

Figure S3 plots 3D and 2D dynamic signatures of the H358 lung cancer cell line and the MDA-MB-231 breast cancer cell line, respectively, under normal culture conditions. Comparing these cell lines with MIAPaCa-2 cells, we found that different cell lines have different dynamic signatures. The value of *length/speed* is proportional to the number of imaging frames that the LD trajectories occupy. Two populations are identified in the MIAPaCa-2 *length*-*speed* graph (Figure S4A). The LDs appeared in higher frame numbers (red gate in Figure S4A) are less likely to move out of the focal plane and tend to have a slower speed (Figure 1K, and Figure S4). The evidence suggests that LDs forming this population are more likely associated with ER and lipid synthesis. Such a conclusion was further verified by comparing the 2D dynamic signatures in the *length-speed* domain (Figure S5) during starvation which is known to reduce LD synthesis and increase LD degradation. H358 cells have a higher percentage of LDs associated with this population (Figure S3C), implying a different LD metabolic rate in H358 cells compared to MIAPaCa-2 cells.

We can gate subpopulations of the trajectories in the *DF-maxd* domain and further explore the *speed* and *length* features. We gated the trajectories having *DF* >10 (red) and *maxd* >3 (blue) (Figure 1J) and plotted the speed and length values for both populations. The results indicate that less directional LDs tend to have a lower *speed* and a shorter *length* (Figures 1K,L). These features unveil the dynamic characteristics of different types of LDs in cells.

The quantification methods developed here allow us to investigate LD dynamics and cellular lipid metabolism from a novel angle. We subsequently explored the responses of LD dynamics of living cells in response to various stimuli.

### Staining perturbations to LD dynamics

Lipophilic dyes are commonly used to visualize LDs in living cells. We compared LD dynamic signatures using our hybrid CARS-TPEF microscope for living cells with and without fluorescent labeling. BODIPY green was used to label LDs in MIAPaCa-2 cells and CARS/TPEF time-lapse image stacks were acquired to quantify the LD dynamics. Compared to the non-labeled control group, BODIPY labeling groups with 2 µg/mL and 4 µg/mL dye concentration for 1 hr, as well as with 2 µg/mL for 6 hr, show clear shifts of *DF* towards higher values and shifts of *maxd* towards lower values (Figure S6). We fit the *maxd* and *DF* values for all four cases (Figure S7A,C) and compared the derived *Xc* value which is associated with the median value of the curves. The results clearly show decreased *Xc* values for two of the labeled groups in the *maxd* domain and increased *Xc* values for all labeled groups in the *DF* domain (Figure S7B,D). A 2D plot of *Xc* values in *DF* and *maxd* domains of all four cases allows us to better distinguish the staining-induced change of LD dynamics (Figure S7E). Further, we can take the ratio of LDs having *DF* > 10 and *maxd* > 3 μm, and the ratio of LDs having *maxd* < 2 μm and *DF* < 5, both of which show clear increases for the all three labeled cases (Figure S7F). These changes indicate LDs tend to reduce their movement, directionality, and relocation distance when BODIPY is used. This cellular perturbation suggests that label-free imaging using CARS microscopy is a more appropriate way to quantify LD dynamics than using fluorescent dyes.

From the CARS images, we discovered that the 6 hr treatment of MIAPaCa-2 cells with 2 µg/mL BODIPY can increase cell membrane blebbing (Figure S8), and 1 hr treatment of MIAPaCa-2 cells with 4 µg/mL can change the morphology of the cells to a more rounded shape (Figure S9), both of which confirmed the functional perturbation of the BODIPY dye to live cells.

### Low-temperature effect on LD dynamics

In addition to the perturbation due to staining, another common environmental stimulus for live cells is a change in temperature. It is believed that cell metabolism scales with temperature because the enzyme activity is only optimal within a relatively small temperature range. We explore how LD dynamics change with temperature, offering a label-free indicator of cellular response due to temperature change. We first measured the MIAPaCa-2 LD dynamics at 37 °C. Then we turned off the temperature maintaining system and exposed the live cells to room temperature (24 °C) for 1 hr before measurement. Next, we reheated the sample back to 37 °C and measured the recovering process. The dynamic signature projected onto the *DF-maxd* domain for the three conditions are shown in Figures 2A-C. We found that the 1 hr 24 °C exposure tends to shift the LDs towards a higher *DF* value and a lower *maxd* value, indicating reduced active transportation and increased non-directional movement impacted by the hypothermia exposure. The *maxd* and *DF* histograms of LDs in all conditions, together with the lognormal fitting results, are shown as Figures 2D,F, respectively. We derive the median-value-associated *Xc* values from the lognormal fitting, and the *σ* values, which represent the skewness of the curves. A comparison of these values (Figures 2E,G) shows that both the median values and the skewness of *maxd* histograms decreased after 1 hr of 24 °C exposure, and recovered to the 37 °C levels when the samples were reheated back to 37 °C. These results indicate that short-term hypothermia exposure of these living cells does not cause irreversible metabolic changes.

**Figure 2.**
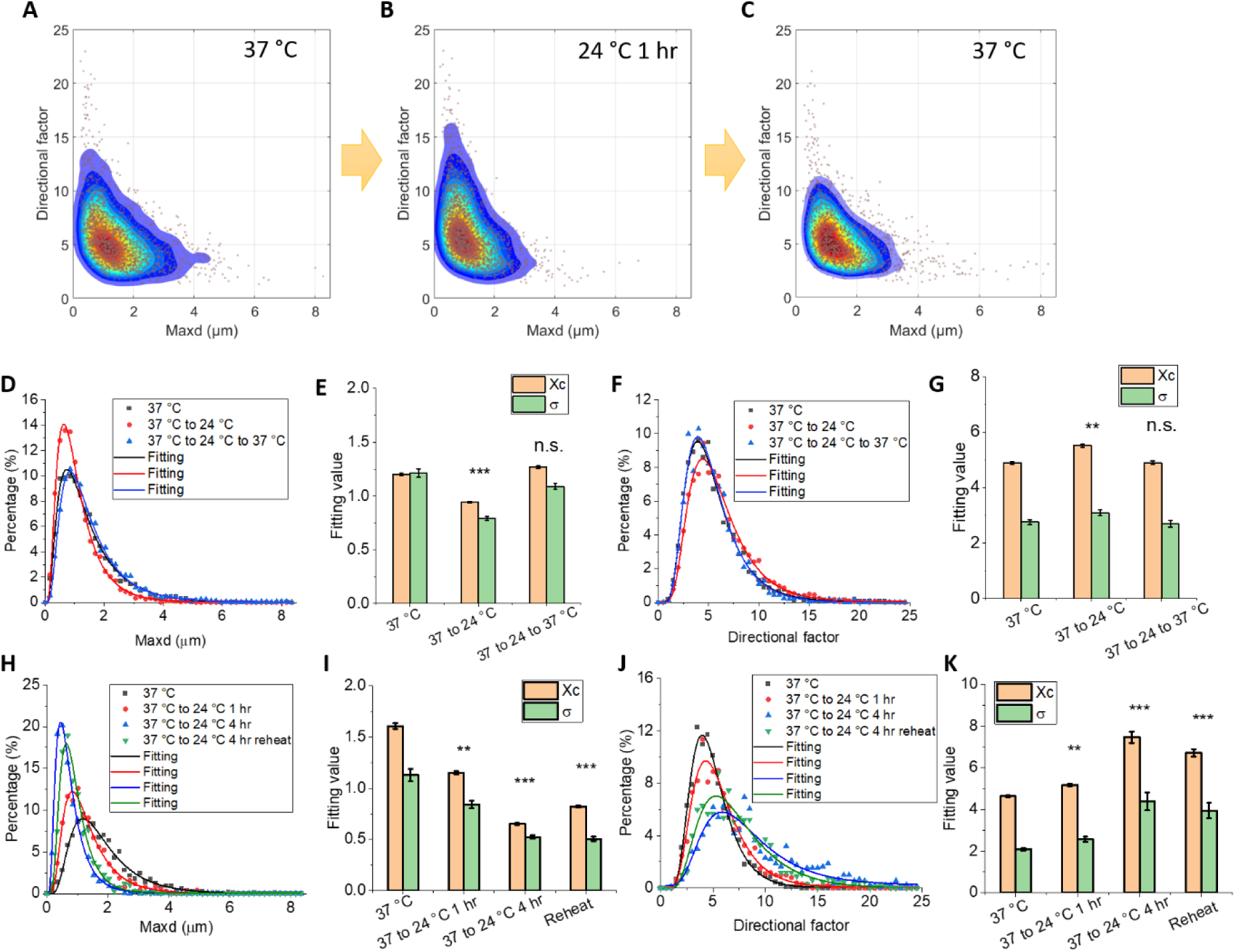
Detection and quantification of hypothermia-induced changes in LD dynamics. (A)-(C) Dynamic signatures of MIAPaCa-2 cells at 37 °C, hypothermia exposure to 24 °C for 1 hr, and reheating of sample to 37 °C, respectively, projected onto the *DF-maxd* plane. (D) Histograms (dots) and the lognormal fitting of the histograms (solid curves) of MIAPaCa-2 cells in the *maxd* domain from the control (37 °C, black), 1 hr hypothermia exposure (24 °C, red), and after reheating (37 °C, blue) conditions. (E) *Xc* and *σ* derived from the lognormal fitting from each condition in panel (D). (F) Histograms (dots) and the lognormal fitting of the histograms (solid curves) of MIAPaCa-2 cells in the *DF* domain, from the three conditions. (G) *Xc* and *σ* derived from the lognormal fitting from each condition in panel (F). (H) Histograms (dots) and the lognormal fitting of the histograms (solid curves) of MIAPaCa-2 cells in the *maxd* domain from the control (37 °C, black), 1 hr hypothermia exposure (24 °C, red), 4 hr hypothermia exposure (24 °C, blue), and after reheating (37 °C, green) conditions. (I) *Xc* and *σ* derived from the lognormal fitting from each condition in panel (H). (J) Histograms (dots) and the lognormal fitting of the histograms (solid curves) of MIAPaCa-2 cells in the *DF* domain from the four conditions in panel (H). (K) *Xc* and *σ* derived from the lognormal fitting from each condition in panel (J). For the student’s t-tests for the 1 hr hypothermia exposure, n=9, and the 4 hr hypothermia exposure, n=7.

Next, we studied the effect of longer hypothermia exposure. The MIAPaCa-2 cells were maintained at 24 °C for up to 4 hr. We measured LD dynamics before hypothermia, at 1 hr exposure, 4 hr exposure, and after the samples were reheated back to 37 °C after the 4 hr exposure. From the *DF-maxd* domain plots, as shown in Figure S10, we discovered continuous shifts towards larger *DF* and smaller *maxd* values as a function of hypothermia exposure time. The fitting results reveal a decrease in both the *Xc* and the *σ* on the *maxd* domain (Figures 2H,I), and an increase of these values on the *DF* domain (Figures 2J,K) induced by the low temperature. Interestingly, we found that after reheating the sample back to 37 °C, the LD dynamic signatures did not recover to their original levels (Figure S10 and Figures 2H-K). These results indicate that a 4 hr hypothermia exposure causes irreversible changes to cell functions or the changes that cannot be reversed in the first hour of recovery.

### Drug-induced apoptosis

Apoptosis is a common and highly regulated process controlling cell death. Analyzing the LD movement, we can better understand this process from a dynamic angle and identify the dynamic signatures indicative of this change. To induce apoptosis, we treated cells with staurosporine (STS) (500 nM final concentration) and measured the LD dynamics as a function of treatment time (0, 1, 3, 11 hr). From the *DF-maxd* plots, we found that STS treatment tends to shift the LD trajectories towards larger *DF* and smaller *maxd* values (Figures 3A-D). Such a change was amplified for longer STS treatment. Histogram fitting in the *maxd* domain quantifies the continuous decrease in both the median value and the skewness of LD trafficking distribution induced by STS and treatment time (Figures 3E,F), which can be correlated in the *Xc-σ* space (Figure 3G). Similarly, the analysis in the *DF* domain shows increases in the median value and the skewness of LD trafficking as a function of treatment time (Figures 3H-J).

**Figure 3.**
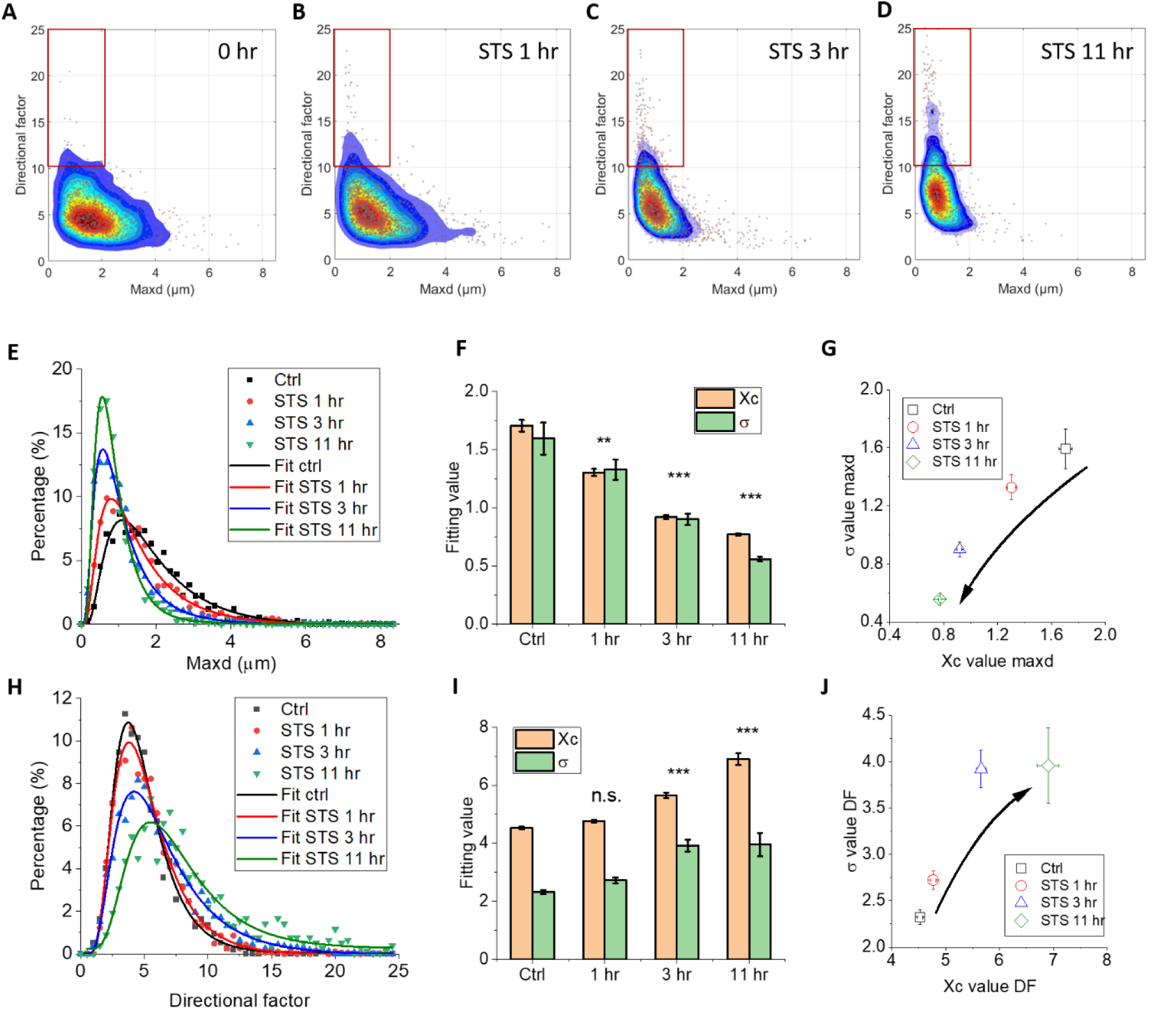
Changes in LD dynamics during apoptosis. (A)-(D) Dynamic signatures of MIAPaCa-2 control group, 0.05% STS treatment for 1 hr, 3 hr, and 11 hr, respectively, projected onto the *DF-maxd* plane. The red rectangles are shown as reference marks. (E) Histograms (dots) and the lognormal fitting of the histograms (solid curves) of MIAPaCa-2 cells in the *maxd* domain from the control (black), 1 hr STS treatment (red), 3 hr STS treatment (blue), and 11 hr STS treatment (green). (F) *Xc* and *σ* values derived from the lognormal fitting from each condition in panel (E). (G) Time-dependent STS treatment results plotted in the *maxd Xc-σ* domain. The arrow indicates the change over STS treatment time. (H) Histograms (dots) and the lognormal fitting of the histograms (solid curves) of MIAPaCa-2 cells in the *DF* domain from the four conditions. (I) *Xc* and *σ* values derived from the lognormal fitting of each condition in panel (G). (J) Time-dependent STS treatment results plotted in the *DF Xc-σ* domain. The arrow indicates the change over STS treatment time. For the student’s t-tests, n=6.

Such LD dynamic changes were reproduced by using a higher concentration of STS (1 μM). Very similar trends were observed with much-amplified changes at a similar treatment period (Figure S11). The major biological function of STS is binding to protein kinase to prevent adenosine triphosphate (ATP) binding to the same enzyme. It was reported that STS can produce a time-dependent and concentration-dependent increase in the caspase-3 activity, which is linked to DNA fragmentation (Belmokhtar et al., 2001; Yue et al., 1998). Studies also showed that STS can alter the mitochondrial membrane potential and causes cytochrome c release (Ahlemeyer et al., 2002; Mirkes and Little, 2000). One of the possible causes of changes in LD dynamics is likely due to the reduction of ATP production because of the STS-induced mitochondria dysfunction. The active transport of LDs, which is usually more directional and with a larger *maxd* value, is powered by ATP.

We used MIAPaCa-2 cells to perform the majority of the studies. As demonstrated in Figure S3, different cells might have different dynamic signatures. However, they might share similar changes in certain conditions. For example, we compared changes of LD dynamics of H358 cells during hypothermia exposure and starvation (Figure S12) and found very similar trends as the MIAPaCa-2 cells (Figures 2,4). If we compare the dynamic signature changes in this apoptosis study with the previous fluorescent labeling and temperature-dependent studies, we find that all perturbations tend to induce decreases in the directionality and the *maxd* for LD movements, which are likely common signatures for perturbed, unhealthy, and ‘unhappy’ cells.

**Figure 4.**
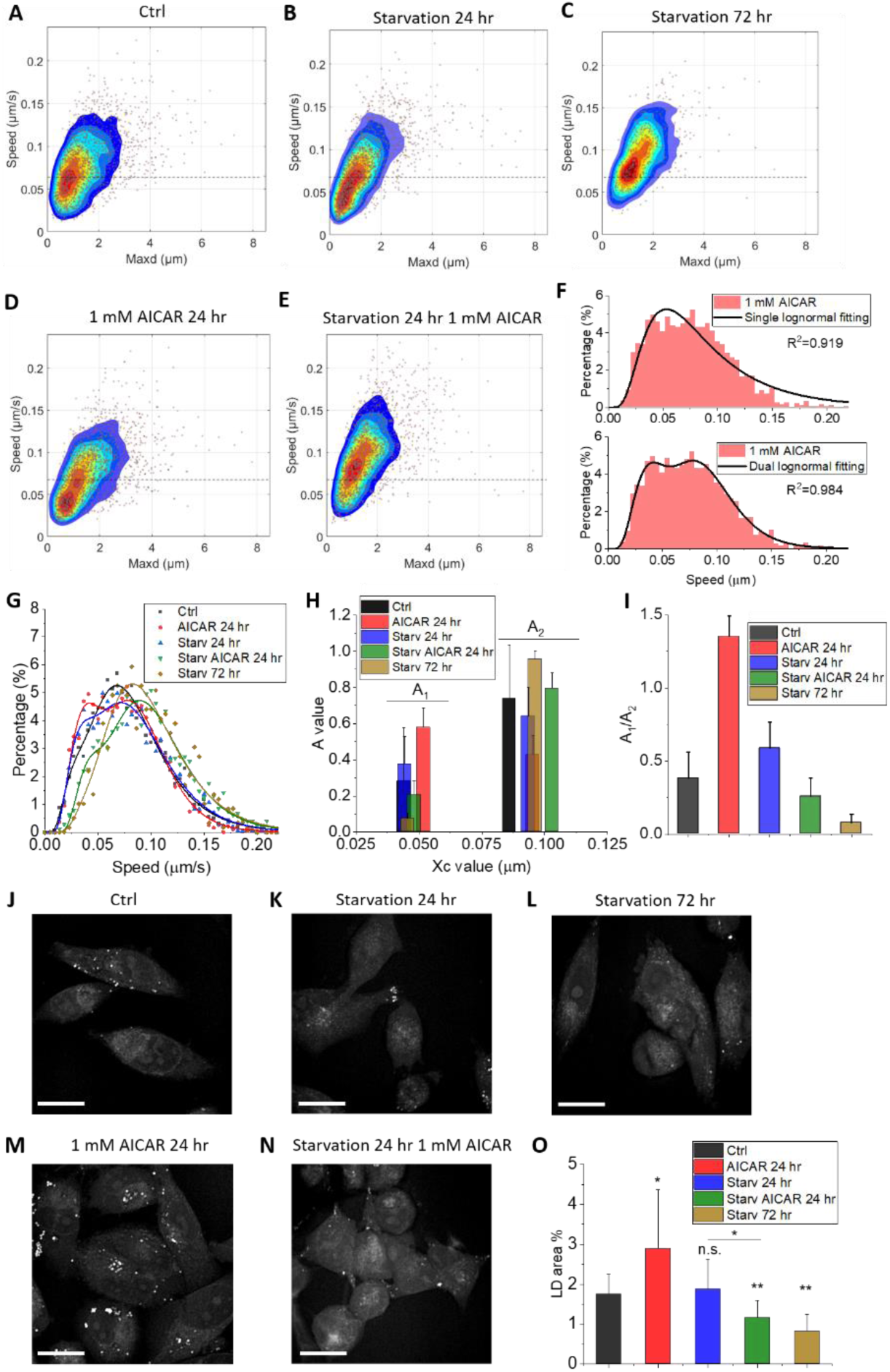
Changes in LD dynamics related to AMPK activation. (A)-(E) Dynamic signatures of the MIAPaCa-2 control, 24 hr starvation ((-)glucose, (-)FBS), 72 hr starvation, 1 mM AICAR treatment in normal culture medium for 24 hr, and 1 mM AICAR treatment in starvation condition for 24 hr, projected onto the *speed-maxd* plane, respectively. The dotted lines are shown as reference marks. (F) Comparing the fitting of *speed* histogram of MIAPaCa-2 cells using a single lognormal distribution function (upper panel) and a dual lognormal distribution function (lower panel). (G) Histograms (dots) and the dual-lognormal fitting of the histograms (solid curves) of the dynamics signatures in the *speed* domain for the five conditions in panels (A-E). (H) *Xc* and amplitude values reflect the median *speed* and the population fractions of the two subpopulations of LDs in MIAPaCa-2 cells in the five conditions. (I) The ratio of amplitudes A_1_/A_2_ for the five conditions. (J-N) Sample CARS images from the five conditions in greyscale. (O) Quantification of LD area percentage for the five conditions using the method described in Figure S15. For the student’s t-tests, n=8.

### Starvation and AMPK activation

LDs are energy storage reservoirs and regulators in cells. It is therefore critical to understand how energy metabolism is associated with LD dynamics, and if the LD trafficking information can provide additional insights for cell metabolism. We explored metabolic manipulations related to the AMPK pathway, a central signaling pathway of energy homeostasis of cells (Hardie et al., 2012). AMPK pathway can be activated through energy starvation or small activator molecules (Hardie, 2016; Kim et al., 2016; SALT et al., 1998). We first created four conditions to explore the effects of glucose and fatty acids on LD dynamics, including control ((+)glucose, (+)fatty acids), glucose only ((+)glucose, (-)fatty acids), FBS only ((-)glucose, (+)fatty acids), and no glucose no FBS ((-)glucose, (-)fatty acids). MIAPaCa-2 cells were cultured under these conditions and measured at 24 hr and 72 hr time points. We plotted the dynamic signatures of MIAPaCa-2 cells in the *speed-maxd* domain for all the conditions (Figure S13) and found an increase in the overall speed at 72 hr but not at 24 hr starvation, with both glucose and fatty acids deprived (Figures 4A-C). Then we studied the same cells treated with 1 mM 5-aminoimidazole-4-carboxamide-1-β-D-ribofuranoside (AICAR), an AMPK agonist and direct activator, in normal and starved culture conditions, and found a similar transition of the dynamic signature in the latter but not in the former condition (Figures 4D,E). Since both starvation and AMPK activation activates the AMPK pathway, which effectively slows down fatty acid synthesis and increases degradation, the very similar trends in Figures 4C,E are likely indicators of AMPK activation. AICAR treatment speeds up such a change in the starved condition since 24 hr starvation by itself does not cause the same change (Figure 4B).

AMPK activation causes reduced LD synthesis and increased LD degradation. Therefore, fewer LDs are associated with the ER, and more are transitioned to the cytosol for degradation and active transportation. From previous studies, we discovered that active transportation tends to have higher *speed* and *maxd* values compared to the ER-associated non-directional movement (see Figures 1J,K). Consequently, the metabolic signature change in the *speed-maxd* domain associated with AMPK activation is the increase in the *speed* and *maxd* values.

If we compare the AICAR treatment group under the normal condition (Figure 4D) with the non-treated group (Figure 4A), we found a decrease in the LD speed. This indicates AICAR treatment of MIAPaCa-2 cells without starvation is causing an opposite change to LD dynamics compared to the AMPK activation. Such a change implies an increased LD synthesis/degradation ratio. MTT cell proliferation assay results show that MIAPaCa-2 cells are experiencing an increased proliferation after being treated by the AICAR in normal culture media (Figure S14). This indicates the continued growth of the cells, which can be correlated with the increased LD synthesis.

To further understand the changes in LD dynamics, we plot the histograms of LD trajectories on the *speed* axis for quantitative comparison. We found that a single lognormal distribution function cannot achieve good fitting of the histogram (Figure 4F, upper panel). This is because subpopulations of LDs in the cells are better separated in the *speed* domain. Using a dual-lognormal function, we can better fit the *speed* histogram (Figure 4F, bottom panel) and distinguish the two LD populations having different *speed* distributions. We fit the MIAPaCa-2 *speed* histogram for all the five conditions (Figure 4G) and plot the amplitude (A) values, which represent the fractions of the LDs in each population, over *Xc*, which associates with the median *speed* of the population (Figure 4H). We found that for all conditions, there are two pools of LDs, one (A_1_) centered around 0.05 μm/s with the other (A_2_) around 0.1 μm/s. A_1_ is the population having small *speed* and *maxd* values, thus associated with ER and LD synthesis; while A_2_ is the population having larger *speed* and *maxd* values, thus associated with cytosol and LD degradation. If we take the A_1_/A_2_ ratio, we find that AICAR treatment under normal conditions has significantly increased this ratio compared to the control, that 24 hr starvation has a similar ratio as the control, while AICAR + starvation for 24 hr and starvation for 72 hr have much-reduced ratios (Figure 4I). These changes reveal the differences in the ratio between the two pools of LDs under different conditions.

We then analyzed the quantity of LDs using CARS images taken from the five conditions. Sample images are shown as Figures 4J-N, from which we can see an obvious increase in the total LD amount for the AICAR treatment group under the normal condition (Figure 4M). We quantify the total LD amount using an intensity thresholding method as displayed in Figure S15, and found that the LD area percentage of the cells for the five conditions has very similar trends as the A_1_/A_2_ ratio we measured from the dynamic analysis (Figures 4I,O). These results indicate that an increased ratio of LDs in the A_1_ group which is associated with LD synthesis is directly linked to a higher amount of total LDs in cells, suggesting a higher rate of LD synthesis which is observed in the AICAR+normal condition group. On the other hand, higher rates of LD degradation and decreased total LD amount are detected from the AICAR+starvation and 72 hr starvation groups, likely caused by the AMPK activation.

If we look at the 24 hr starvation group with only FBS (Figure S13D), we find a decrease of LD speed. We quantify the A_1_ and A_2_ LD groups by fitting the *speed* histograms in this condition and compare the results with the control group. We discovered an increase in the A_1_/A_2_ ratio, which indicates increased LD synthesis (Figure S16A-F). From these results, we predict that compared to the control group, fatty acids alone as the energy source would increase the LD synthesis. This conclusion was further confirmed by the analysis of the LD amount using the intensity thresholding method discussed before (Figure S16G-I).

### Lipid-metabolism-related enzyme inhibition

After exploring the AMPK-related effects, we directly perturbed the lipid metabolism by targeting different fatty-acid-regulating enzymes and studied the LD dynamics associated with these perturbations. Two compounds were used to inhibit fatty acid degradation: ranolazine, which inhibits thiolase (Kantor et al., 2000; Sabbah et al., 2002), and etomoxir, which inhibits carnitine palmitoyltransferase I (CPTI) (Reaven et al., 1988). One chemical, C75, was used to inhibit fatty acid synthase (FASN) (Kim et al., 2004). Besides, tunicamycin was deployed to induce ER stress (Banerjee et al., 2011). From the dynamic signatures in the *speed-maxd* domain, we found that the LD *speed* values of MIAPaCa-2 cells treated by fatty acid degradation inhibitors were much lower than those treated by the ER-stress inducer and the FASN inhibitor (Figures 5A-D). Quantitative histogram fitting on the *speed* domain provides A_1_ and A_2_ population information for all four treatment groups (Figure 5E). Comparing with the control group, the A_1_/A_2_ population ratio (Figure 5F) shows an increased fraction of synthesis-related LDs for thiolase inhibition (ranolazine), similar fractions of synthesis-related LDs for CPT1 inhibition (etomoxir), and reduced fractions of synthesis-related LDs for ER stress (tunicamycin) and FASN inhibition (C75). We also compared the *maxd* values for all the conditions and found that the *Xc* values of *maxd* follow the opposite trend as the A_1_/A_2_ ratio (Figure 5G). Since A_1_ is associated with LD synthesis and A_2_ is linked to LD degradation, while LD synthesis is correlated with non-directional movement on ER and LD degradation is linked to active transportation, the trends in *maxd* reflect a similar metabolic change in cells as was unveiled by the A_1_/A_2_ ratio. We further plotted the A_1_/A_2_ ratio against the *maxd Xc* value on a 2D graph (Figure 5H) and found that increased and reduced LD synthesis processes are diverging to different directions from the control point. Sample CARS images from the four treatment groups are shown in Figures 5I-L. The ER areas of the MIAPaCa-2 cells have higher CH_2_ CARS intensities than the cytosol, but lower signal than the LDs. Using intensity thresholding, we can highlight the ER areas from the cells directly from CARS images (Figure S17), which agrees well with the results from ER labeling using ER-Tracker™ Green (Figure S17). CARS images show that the LD-degradation inhibited groups have more ER-bound LDs compared to the ER-stress and the FASN inhibited groups (see green areas in Figures 5I-L).

**Figure 5.**
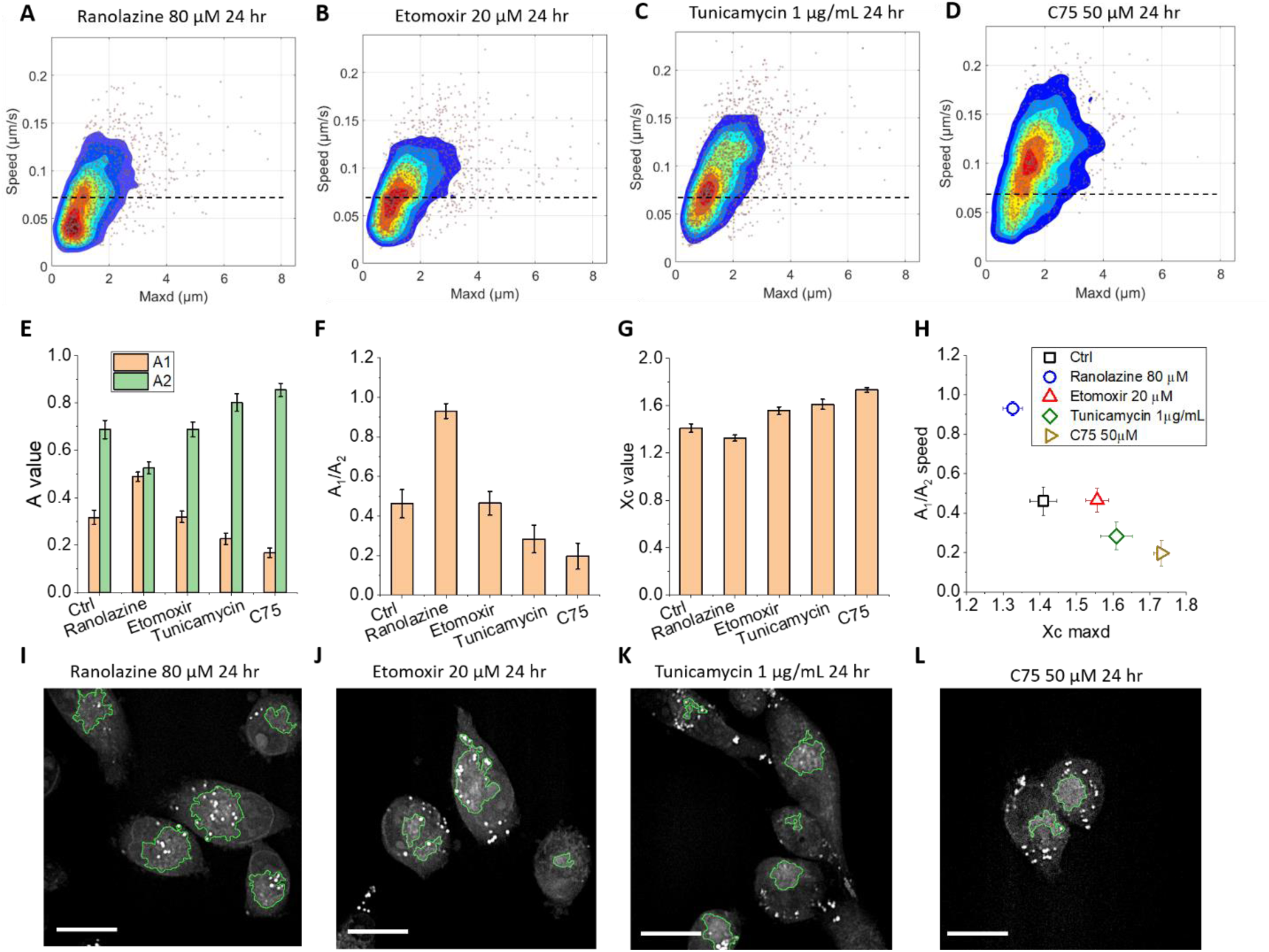
Changes in LD dynamics related to inhibition of lipid metabolic enzymes. (A)-(D) Dynamic signatures of MIAPaCa-2 treated by 80 μM ranolazine, 20 μM etomoxir, 1 μg/mL tunicamycin, and 50 μM C75, projected onto the *speed-maxd* plane, respectively. The dotted lines are shown as reference marks. (E) Amplitudes A_1_ and A_2_ values, reflecting the fractions of two subpopulations of LDs in cells, derived from fitting *speed* histograms using dual-lognormal distribution. (F) The ratio of amplitudes A_1_/A_2_ for the five conditions. (G) The *Xc* values obtained from fitting *maxd* histograms using single-lognormal distribution functions. (H) The control group and four treatment groups plotted on the A_1_/A_2_ – *maxd Xc* domain. (I-L) Sample images of MIAPaCa-2 cells from the four treatment groups. The green area plots the intensity-thresholding-highlighted ER areas. The intensity thresholding method for ER analysis is illustrated in Figure S7.

We also studied the effect of another thiolase inhibitor, trimetazidine (Figure S18) (Kantor et al., 2000). Interestingly, from the histogram fitting (Figure S18B,C), we found this case does not fall onto the curve of other conditions in the A_1_/A_2_ - *maxd Xc* space (Figure S18D), but deviates from the control toward a new direction. Such an abnormal change in LD dynamics indicates something abnormal happened for cell metabolism. We investigated the CARS images and found that trimetazidine tends to induce LD aggregation, as shown in Figure S18E. Many LDs aggregated after 24 hr treatment and did not separate even over 220 s (Figure S18E and Video S2). These LD aggregates are in the cytosol, resulting in a low A_1_/A_2_ ratio. Since different LDs are linked to different microtubules and are unlikely to move long distances once aggregated, these assemblies have much less *maxd* value, bringing down the median value (*Xc*) of the entire *maxd* histogram.

## Discussion

For living cells, if an image is their ‘ID card’, then their dynamics are ‘live-videos’ of their behaviors. Dynamics provide useful information that cannot be derived from static images. Through the quantification of LD dynamics in real-time using a label-free approach, we derived the speed, displacement, length, and directionality of LD movements in cells, which allowed us to monitor environmental- and stimuli-induced changes in cellular dynamics in real-time. We, for the first time, quantify the dynamic changes by histogram fitting and comparative analysis. Quantification of LD dynamics offers the opportunity to separate different populations of LDs based on their different metabolic processes (LD synthesis and degradation) within cells, and shed new light on the study of lipid metabolism in living cells. The quantitative LD dynamic information, together with label-free imaging, provides scientists with a new method to understand stimuli- and drug-induced metabolic reprogramming in living cells. We expect that the knowledge obtained from these organelle dynamics will help biologists to better understand cell metabolism under stress conditions and better analyze cell responses to various drug treatments.

Several factors impact organelle dynamics in living cells. Reduced LD displacement after short-term hypothermia exposure is most likely induced by the reduction of energy flux, while long-term hypothermia exposure might cause manifold changes in cell metabolism. The impact of STS on LD dynamics is also likely multifarious, including changes in the cellular cytoskeleton, ATP production, and DNA fragmentation. More focused studies are expected to further understand such influences and impacts. Regarding the AMPK activation and lipid metabolism studies, we have observed clear trends in various conditions that were revealed by LD dynamic analysis and can be explained through the changes in the ratio of LD synthesis/degradation. The conclusions obtained from the dynamic analysis also well-correlate with the evidence observed in conventional image-based quantitative analysis. We have therefore extended our understanding of cell metabolism from this novel angle.

We want to point out that there are errors in trajectory tracking and linking, as well as possible false LD particles detected in the experiments. We have used several approaches to minimize the impact of these errors on our conclusions. First, we set a frame number filter to reject trajectories of less than 20 frames. This filtered out the majority of false trajectories caused by intensity variations and spikes in images. We also optimized the particle linking parameters and used the optimal values for the cutoff, link range, and displacement. We noticed that LD dynamic signatures of the same cell line in the same environment might also be different in different days, and therefore measured a control group for each study. The majority of the control groups have quite similar dynamic signatures, though a few of them are slightly different.

Besides the dynamics of LDs, other organelles dynamics might give us more metabolic information. For example, the mitochondrion, which contains a large number of NADH molecules, can be visualized through autofluorescence at 400-500 nm using the TPEF modality. Therefore, CARS/TPEF dual-channel imaging can simultaneously quantify the dynamics of both LDs and mitochondria. To generate detectable TPEF signals from NADH molecules in mitochondria, femtosecond laser pulses, instead of picosecond pulses as used in the current work, are needed. Whether the femtosecond pulses will cause an excess amount of photo-toxicity and perturbation to live cells that would alter the dynamics of organelles needs further investigation. This study is beyond the scope of this current work and will be performed in the future.

## Materials and Methods

### Coherent anti-Stokes Raman scattering and multiphoton fluorescence microscopy

Our CARS microscope was constructed based on an upright microscope frame (BX51, Olympus). A dual-output laser system (Chameleon Discovery, Coherent Inc.) was implemented to generate a fixed laser beam at 1040 nm, used as the pump beam, and a tunable laser beam (from 660 nm to 1300 nm), used as the Stokes beam. The pulse width of both laser beams was ∼100 fs. To reduce peak power and photo-toxicity, we used a pair of SF57 glass rods (152.4 mm length for each, Lattice Electro-Optics) to chirp the pulses to ∼1 ps for pump and ∼2 ps for Stokes beams after combining with a dichroic mirror (see Figure S1). The power of the pump and Stokes beams at the sample was 14 mW and 10 mW, respectively. The CARS signal was acquired by a photomultiplier tube (PMT) (H7422-40, Hamamatsu) placed in the transmission direction with a bandpass filter (650/13 nm, FF01-650/13/25, Semrock) to reject the excitation beams and auto-fluorescence signals from the sample. Multiphoton fluorescence signals from BODIPY-labeled lipids were collected by an epi-direction PMT (H7422-40, Hamamatsu) with a collection of dichroic mirrors and filters (785±BW nm, Di02-R785-25×36, Semrock). Two pre-amplifiers (PMT-4V3, Advanced Research Instruments Corp.) were used to amplify the signals from PMT before the data acquisition system (PCIe-6351, National Instruments). The water immersion objective lens used in this work (LUMPLFLN, Olympus) had a magnification of 40X and a numerical aperture of 0.8.

### Image acquisition and the analysis of LD dynamics

The lab-written image acquisition software was based on LabView. By setting the galvo mirror step voltage as 0.003 V, a 400×400-pixel image covered a field-of-view (FOV) area of 85×85 µm^2^. CARS images for LD dynamic analysis containing 400×400 pixels were collected with a pixel dwell time of 10 µs, corresponding to 1.6 s per image. Considering extra edge-pixels on each line which will be removed after the acquisition and the galvo mirror returning process, the imaging speed is 2.2 s/frame. A total of 100 frames were acquired continuously for each FOV to record the LD dynamics. For all the LD dynamic analyses in this work, typically 5-9 FOVs were analyzed for each condition to provide statistical quantification. The image stacks were examined immediately after image collection. The inferior image stacks containing whole-cell movement, stage drift, focus drift, or other issues were excluded from the analysis.

For each FOV, the image stack was saved as a .txt file (40,000×400 pixels), converted to a 100-frame image stack, and analyzed by a Particle Tracker ImageJ Plugin (44) using the following values for all parameters: radius=0, cutoff=3, percentile=0.5-2 (depends on the LD amount in the image), link range=1, displacement=5. The percentile was chosen to ensure the oversampling of LDs in images, i.e. picking up more particles than the LDs. The false trajectories from non-LD detections will be eliminated after applying a 20-frame filter. All trajectories after particle linking were saved in a .txt file for quantitative analysis. Software written in MATLAB was used to quantify the trajectories. The maximum displacement (*maxd*) was defined as the maximum displacement of the LD movement during the time of image acquisition. The LD *speed* equaled the total trajectory *length* divided by the total time. The *directional factor* (*DF*) was defined as the total trajectory *length* divided by the *maxd*, which was used to determine the directionality of the trajectory. The trajectory shown in Figure 1C was generated using the ImageJ Particle Tracker Plugin (the left panel: visualize all trajectories → double click the selected trajectory → save as .png) and plotted using Origin 2020 (the right panel).

The dynamic signature of the cells under a specific condition was plotted in 3D using MATLAB, with the projections on three planes. The contour density graph of the dynamic signature in each plane was plotted using MATLAB with a modified jet color scale by replacing the lowest blue color with white. The plot ranges for different graphs were scaled with maximum values detected throughout the study (*maxd*: −0.5 to 8.5, *speed*: −0.024 to 0.24, *length*: −1 to 25, *DF*: −1 to 25). The level step for the contour plots was 0.00004. Trajectory gating and subpopulation 2D plots were performed using MATLAB. The fitting of LD trajectories in the *DF-maxd* domain using a reciprocal function (see Figure 1J) was carried out by Origin 2020 and plotted by MATLAB.

Histograms of the LD dynamics on each axis were created using the ImageJ histogram function with 50 bins and fixed min and max values (min=0, max=8.5 for *maxd*, 0.24 for *speed*, 25 for *DF* and *length*). The histograms were normalized by dividing the total number of trajectories after 20 frame-filtering, then multiplied by 100%. The histograms were plotted and fit using a single or dual lognormal distribution function in Origin 2020.

The single-lognormal function is defined as

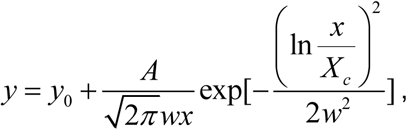

and 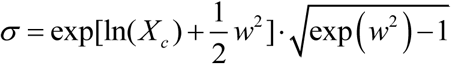

The dual-lognormal function requires self-definition and is defined as

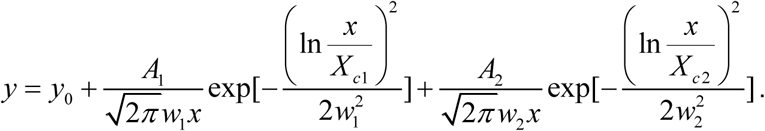

During the fitting, the lower bounds of all parameters were set to 0. The y_0_ intercept was also set to 0. The *Xc* and *σ* values were derived from the fitting for comparison. Dual lognormal distribution was only used to fit the *speed* histogram and gave the A_1_ and A_2_ values roughly equal to the fraction of LDs in each population.

To quantify the significance of changes in LD dynamics, we performed student’s t-tests for LD trajectory information from different conditions. We averaged the measured parameters, e.g. *maxd, speed, length*, and *DF*, for each image stack. Then 5-9 FOVs (n values) were analyzed for each condition to create statistical tables of the parameters. Levels of significance were determined by calculating the p-values comparing the parameter tables from different groups (usually the treatment group v.s. the control group). The asterisks and n.s. have the following meanings: n.s., p > 0.05; *, 0.01 < p < 0.05; **, 0.001 < p < 0.01; ***, p < 0.001.

For all CARS images, a single frame from the .txt image stack file was converted to a .png file for display. For the intensity thresholding, a single image frame was selected for each FOV. We first applied a Gaussian blur function (radius=15) to each image for 10 times (Figure S15B). Then we divided the original image by the Gaussian-blurred image to generate the LD-enhanced image (Figure S15C). This process was used to reduce the impact of background and cell intensity inhomogeneity on the LD particle analysis. From the processed image, we carried out intensity thresholding and used the Particle Analyzer Plugin in ImageJ, selecting particle size < 100 pixels to crop the LD areas (Figure S15F,G). From the original image, we performed intensity thresholding and used particle analyzer to crop the cell area by defining particle sizes > 1000 pixels (Figure S15D,E). Finally, the LD area percentage was calculated as the total LD area divided by the total cell area, multiplied by 100%.

### Temperature control

Except where specifically mentioned, all LD dynamics measurements were performed at 37 °C. The temperature was controlled by a culture dish heating plate (DH-35iL, culture dish incubator, Warner Instruments). To maintain the temperature and reduce media evaporation, we used wet tissue paper to cover the open space between the heating plate and the objective lens. Cells removed from the incubator were immediately placed on the heating plate, unless otherwise mentioned. For hypothermia exposure, the heating plate was turned off and allowed for 5-10 min temperature stabilization to room temperature at 24 °C. The exposure at a given temperature lasted for different periods for different experiments. For reheating, the heating plate was turned back on and 5-10 min was allowed for temperature stabilization at 37 °C. Image collection was started immediately after temperature stabilization.

### Cell culture

MIAPaCa-2, MDA-MB-231, and H358 cells were obtained from ATCC. The cells were normally cultured in DMEM ((+)4.5 g/L glucose, (+)glutamine, (-)sodium pyruvate) mixed with 10% FBS and 1% penicillin-streptomycin (10,000 U/mL). For starvation, glucose-free DMEM was used to eliminate the glucose supply in the medium. The DMEM or glucose-free DMEM with or without FBS created four culture conditions used in the starvation studies, all with 1% antibiotics present. The cells were first seeded to a ∼15-20% confluency followed by 24 hr growth before all the studies or treatments.

### Chemical treatment of living cells

Staurosporine (STS) was obtained from Sigma Aldrich. The Stock solution of STS in dimethyl sulfoxide (DMSO) has a concentration of 1 mM. MIAPaCa-2 cells were treated with STS to reach 500 nM concentrations. Cells were first measured at 37 °C, without any treatment, as the control. Then STS was added to the media to reach the final concentration. CARS image stacks were collected immediately after the treatment, up to one hour. Then the sample was returned to and maintained in the incubator for the later 3 hr and 11 hr measurements. Similar procedures were used for the 1 µM STS treatment.

AICAR treatment of MIAPaCa-2 cells were performed at 1 mM concentration in normal ((+)glucose, (+)FBS) and starved media ((-)glucose, (-)FBS). The treatment time was 24 hr for each condition. Chemicals including ranolazine (80 µM), etomoxir (20 µM), tunicamycin (1 µg/mL), C75 (50 µM), and trimetazidine (1 mM) were used to inhibit different fatty acid-related enzymes or creating ER stress. The concentrations of these chemicals were selected based on those used in the literature to ensure cell viability. All treatments were performed for 24 hr. All the chemicals mentioned above were purchased from Cayman Chemical Company (Ann Arbor, MI) and used immediately after arrival.

### Fluorescent labeling

The BODIPY green (493/503) was purchase from Cayman Chemical Company. A stock solution of 2 mg/mL BODIPY in DMSO was prepared for labeling. Finial concentrations of BODIPY in MIAPaCa-2 cells were 2 µg/mL and 4 µg/mL for different treatments. CARS images and TPEF images were acquired simultaneously from the two channels on the microscope. The LD dynamics analysis was performed using the CARS images, similar to other cases.

ER-Tracker™ Green was purchased from ThermoFisher. The stock solution had a concentration of 1 mM in DMSO. The final concentration of ER-Tracker™ Green for ER labeling was 1 µM and the cells were incubated with the dye for 30 min at 37°C/5% CO_2_ before imaging.

### Cell viability assay

Thiazolyl Blue Tetrazolium Blue (M5655, Sigma Aldrich) (MTT) colorimetric assay was used to measure cell viability. Cells were first seeded in 96-well plates and incubated overnight. Then the inhibitor treatments were performed using AICAR at different concentrations in the normal culture medium. A volume of 100 µL medium was used for each well. Treatment time was 24 hr. The MTT stock solution was prepared at 5 mg/mL in culture medium and was added at one-tenth volume to each well. After 3 hr incubation with the MTT solution, the medium in the wells was replaced by 100 μL of DMSO and shaken well before plate reading. The sampling size was n = 6 for each group. The plate was read by a BD plate reader. The 570 nm absorption was measured for all plates while the 690 nm absorption was also measured as the reference.

## Acknowledgments

The authors thank Aneesh Alex, Jang Hyuk Lee, and Jose Rico-Jimenez from the Center for Optical Molecular Imaging for the initial installation of laser and data acquisition/PC systems, and thank GlaxoSmithKline for the sponsored research support of the Center. We also thank Darold Spillman for lab management and information technology support. This research was financially supported in part by grants from the NIH (R01CA241618, R01EB023232) and the NSF (CBET 18-41539). Additional information can be found at http://biophotonics.illinois.edu.

## Supplemental Information

Supplementary figures and videos are available.

## Supplementary Information for

### Supplementary Figures

**Figure S1.**
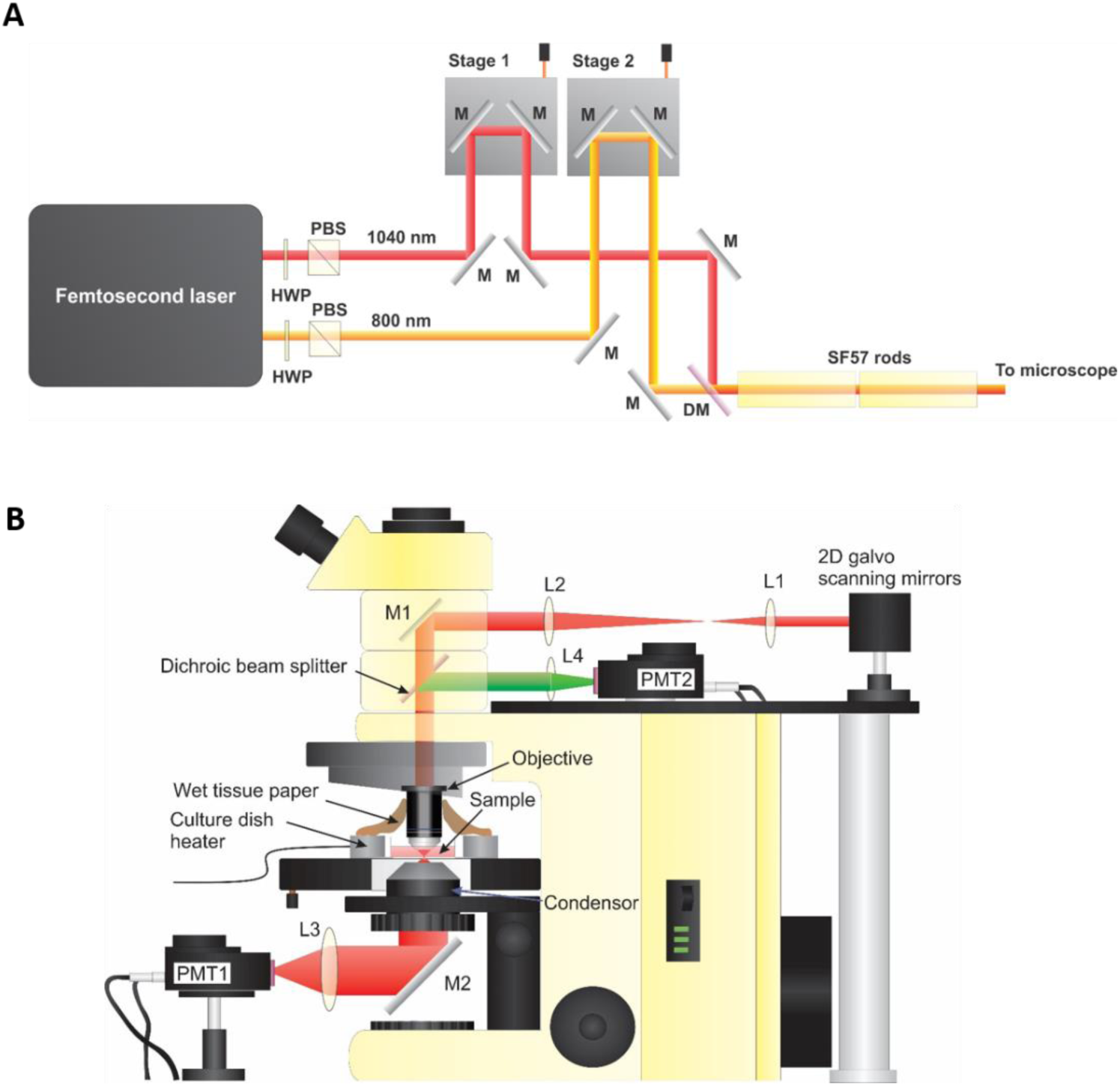
Experimental setup. (A) A schematic showing the coherent anti-Stokes Raman scattering optical paths before the laser-scanning microscope. M: mirrors; PBS: polarization beam splitter; HWP: half-wave-plate; DM: dichroic mirror. (B) A schematic showing the microscope setup used for hybrid-CARS-TPEF imaging. L: lens; M: mirror; PMT: photomultiplier tube.

**Figure S2.**
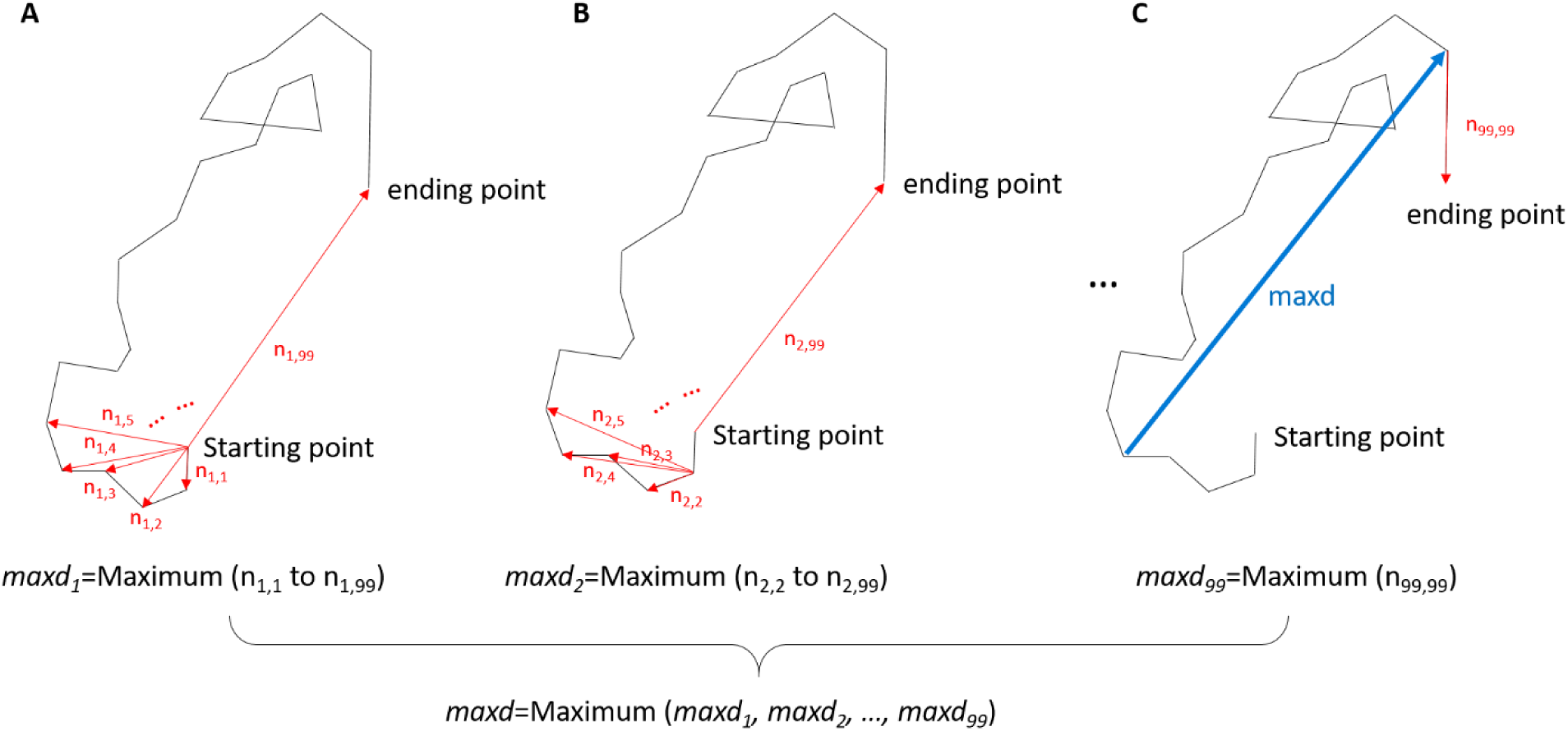
Definition of the maximum displacement. (A) The maximum displacement from the starting point (#1) of a sample LD trajectory. (B) The maximum displacement from the second (#2) time point of the trajectory. (C) The maximum displacement from the second-last (#99) time point of the trajectory, and the definition of the *maxd* of the trajectory.

**Figure S3.**
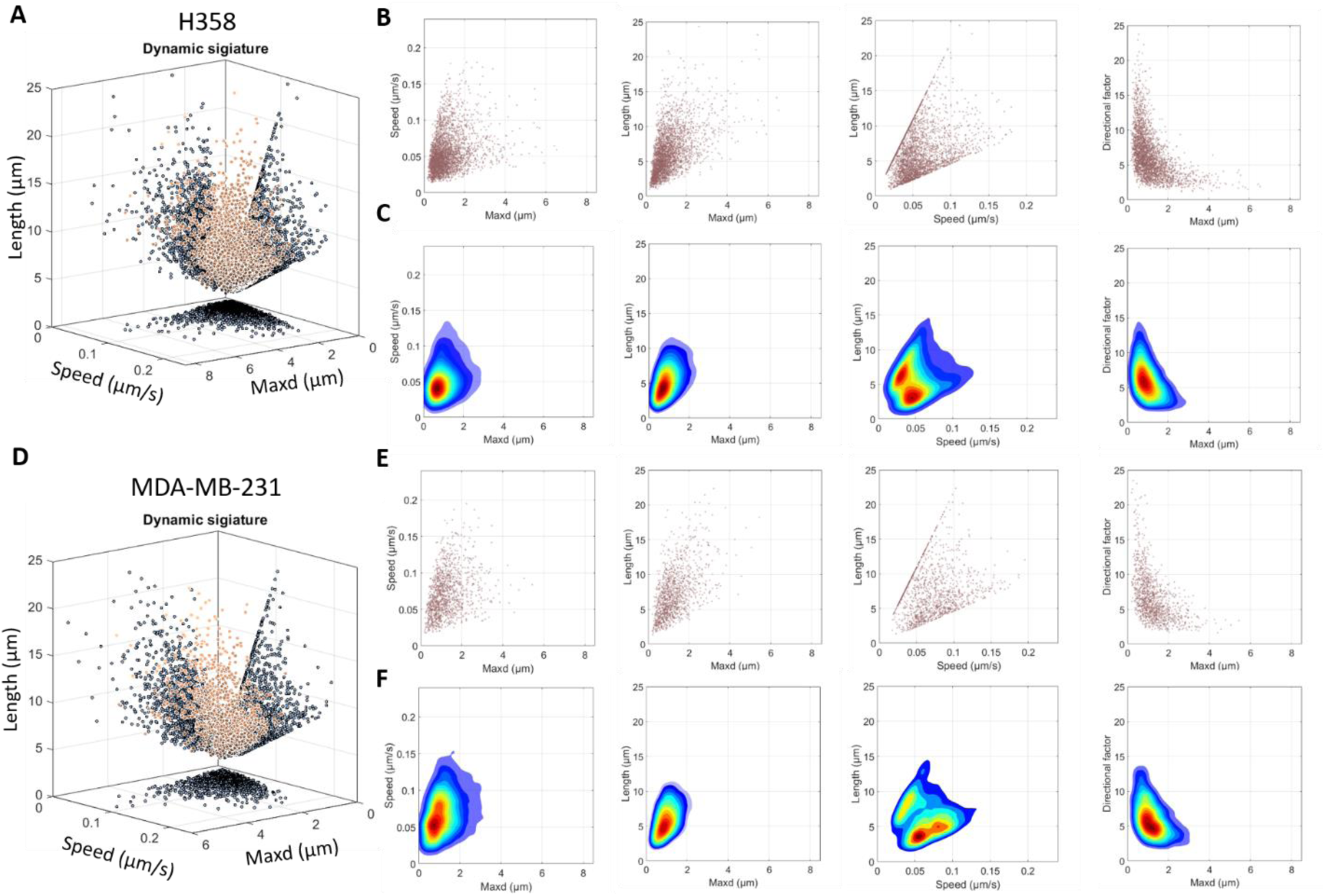
Dynamic signatures of different cell lines. (A) 3D dynamic signature of H358 cells and projections. (B) and (C), scatter and contour plots of H358 dynamic signatures projected onto the *speed-maxd, length-maxd, length-speed*, and *DF-maxd* planes. (D) 3D dynamic signature of MDA-MB-231 cells and projections. (E) and (F), scatter and contour plots of MDA-MB-231 dynamic signatures projected onto the four 2D planes as in panels (B) and (C).

**Figure S4.**
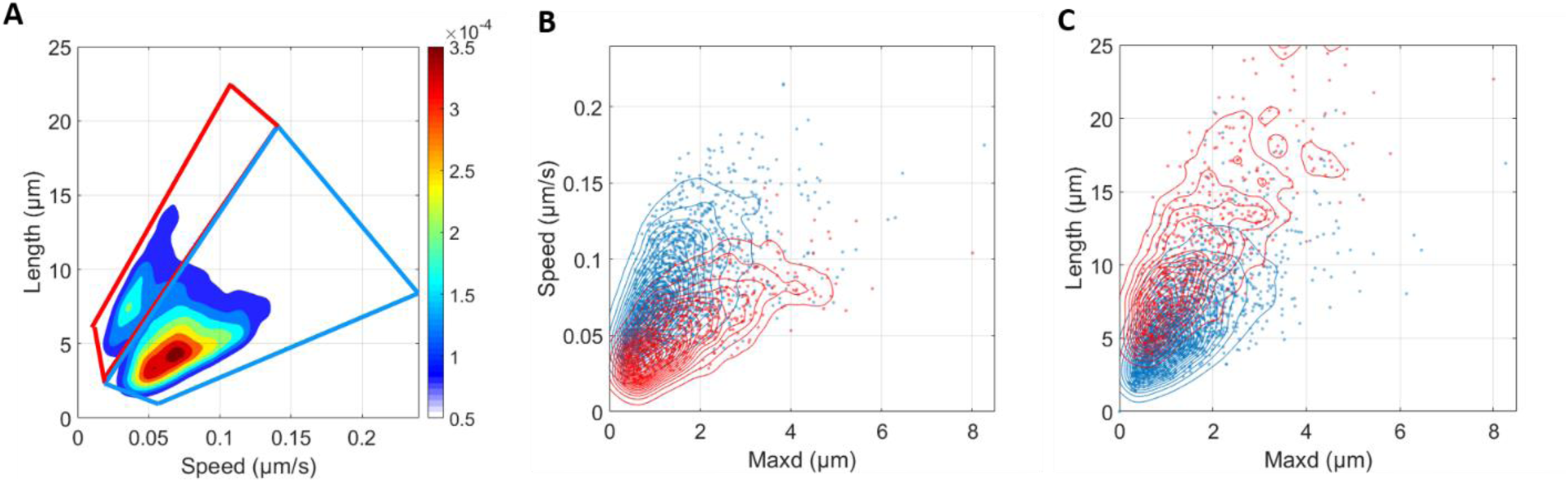
Subpopulation analysis from the length-speed domain. (A) Two gates of the subpopulation of LDs based on the density plot of their trajectories in the *length-speed* domain. (B) and (C) The density plot of the two subpopulations in the *speed-maxd* and the *length-maxd* planes, respectively.

**Figure S5.**
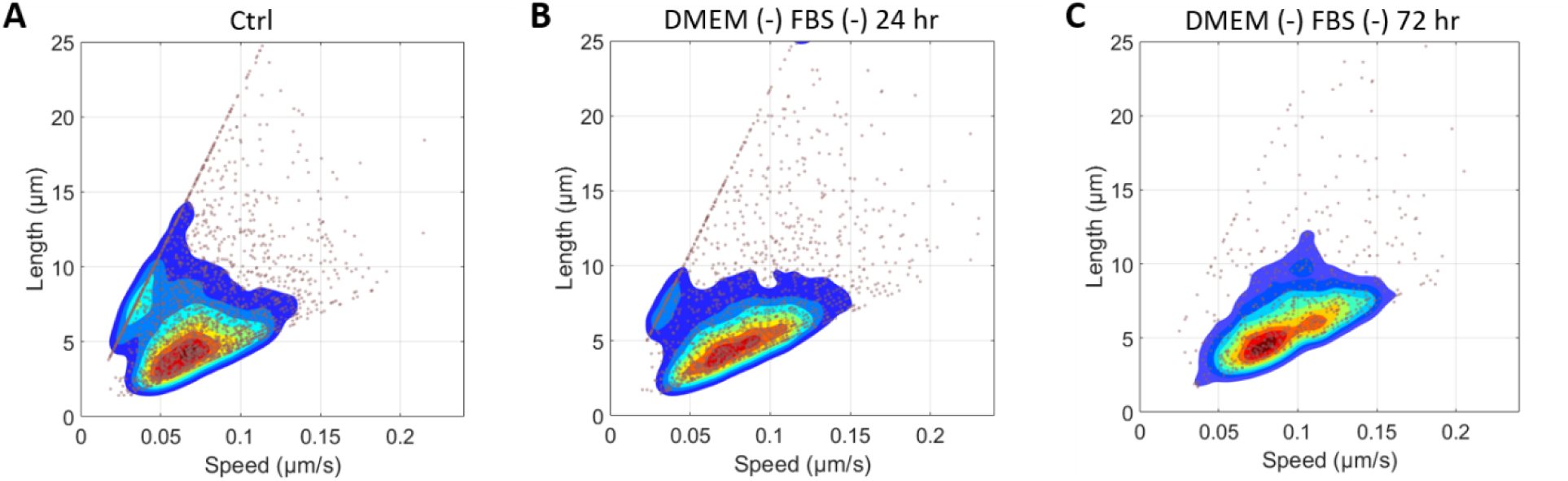
Dynamic signatures of MIAPaCa-2 cells in the length-speed domain in starved conditions. Density plots of the LD trajectories in the (A) control (37 °C, normal culture medium), (B) 24 hr starvation, and (C) 72 hr starvation conditions.

**Figure S6.**
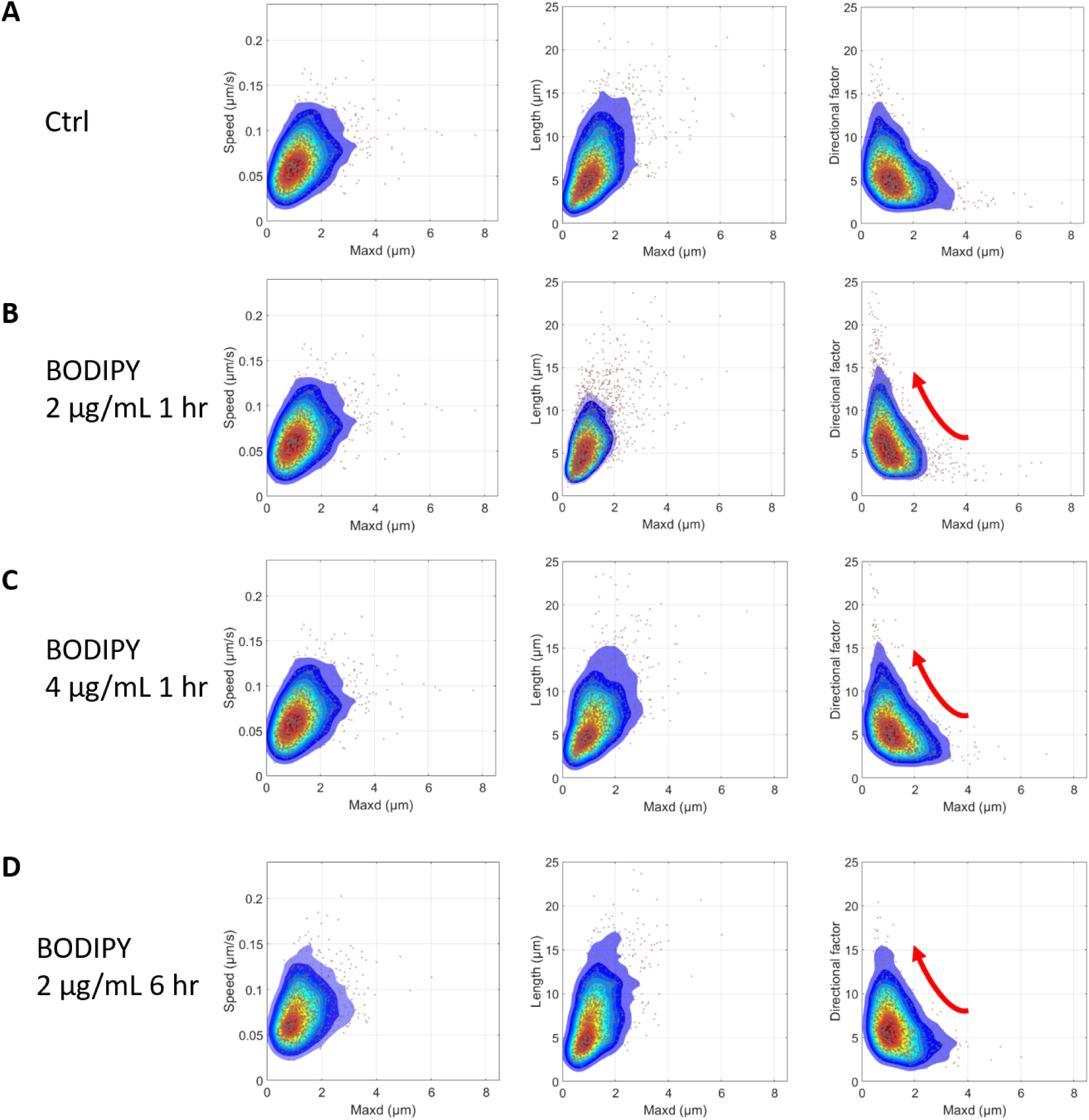
The effects of BODIPY labeling on LD dynamics. (A) The dynamic signature of MIAPaCa-2 cells in the normal condition projected onto the *speed-maxd, length-maxd*, and *DF-maxd* planes. (B)-(D) The dynamic signatures of MIAPaCa-2 treated with 2 µg/mL BODIPY for 1 hr, 4 µg/mL BODIPY for 1 hr, and 2 µg/mL BODIPY for 6 hr projected on the three planes as in panel (A).

**Figure S7.**
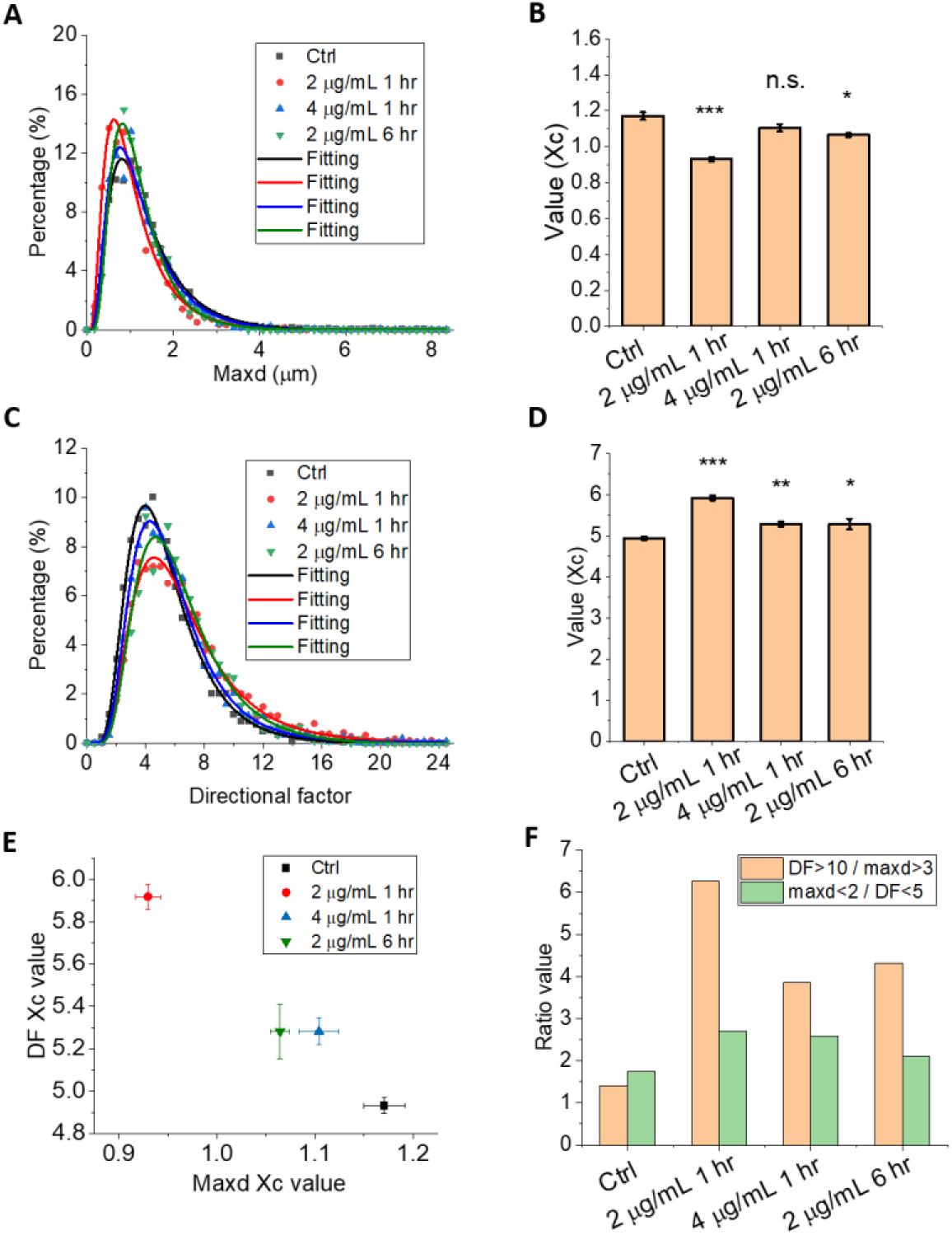
Quantification of the effects of BODIPY labeling on LD dynamics. (A) Histograms (dots) and the lognormal fitting of the histograms (solid curves) of MIAPaCa-2 cells at the *maxd* domain for the control and the three BODIPY treated groups. (B) *Xc* values derived from the histogram fitting in panel (A). (C) Histograms (dots) and the lognormal fitting of the histograms (solid curves) of MIAPaCa-2 cells in the *DF* domain. (D) *Xc* values derived from the histogram fitting in panel (C). (E) *Xc* values of the four conditions plotted on the *DF-maxd* plane. (F) The population ratio of *DF* >10 / *maxd* >3 LDs (yellow) and *maxd* <2 / *DF* <5 LDs (green) for the four experimental conditions.

**Figure S8.**
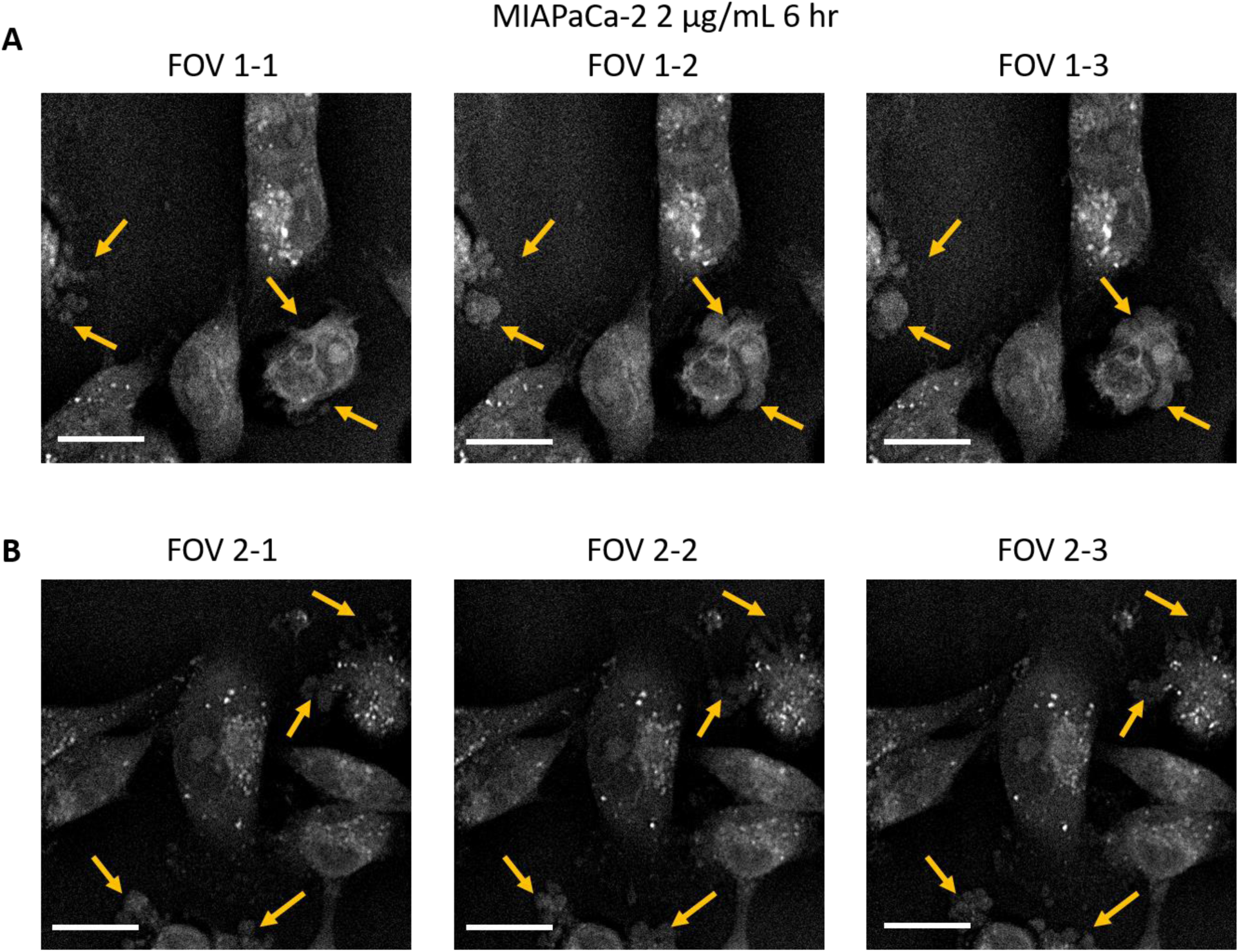
Impact of 2 µg/mL BODIPY on MIAPaCa-2 cells. (A) CARS images from three time-points for MIAPaCa-2 cells labeled with 2 μg/mL BODIPY for 6 hr. The time interval between the two adjacent images is 110 s. (B) Another field of view selected similarly as panel (A). Arrows point out cell membrane blebbing areas. Scale bars: 20 μm.

**Figure S9.**
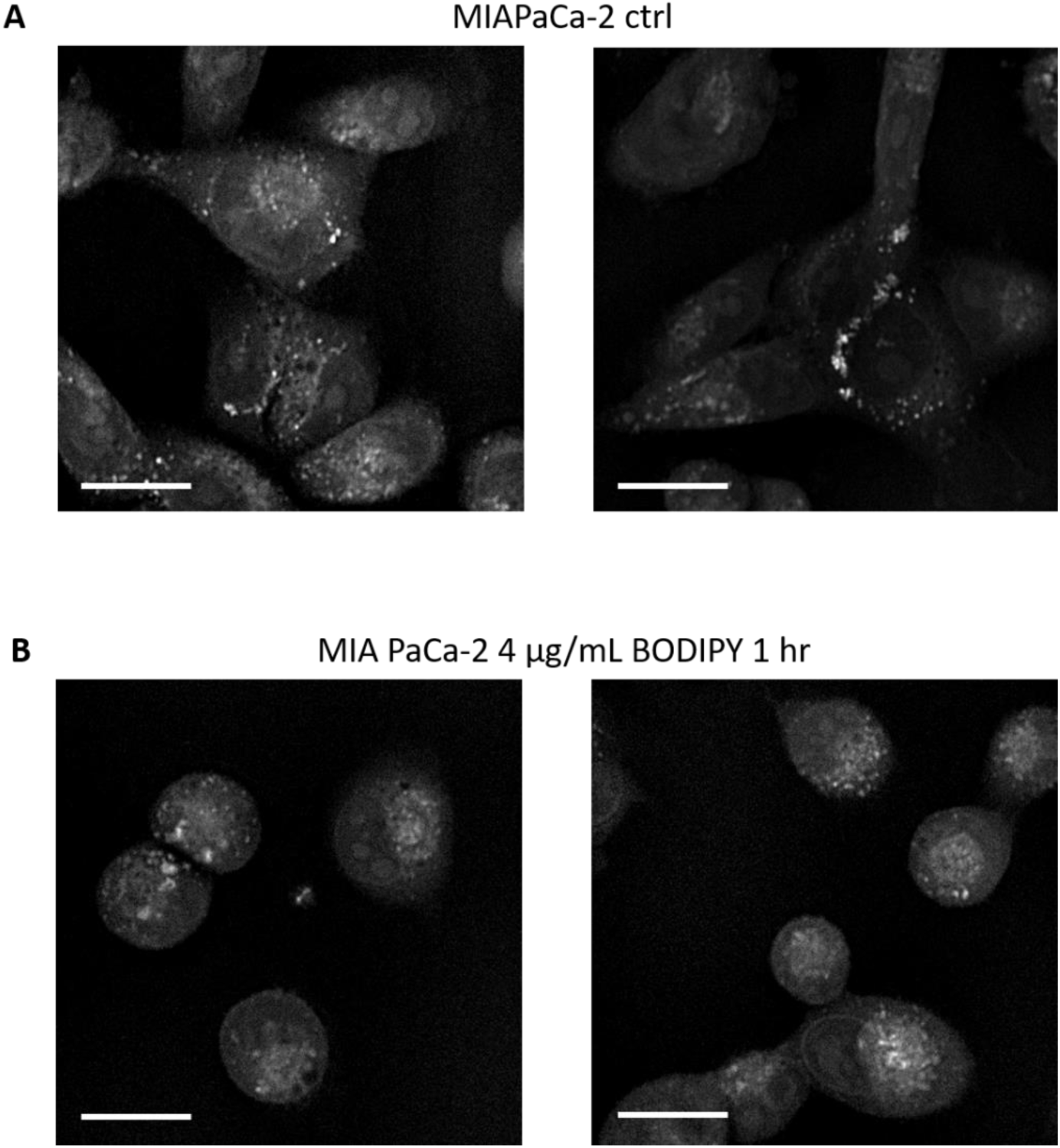
Impact of 4 µg/mL BODIPY on MIAPaCa-2 cells. (A) CARS images from MIAPaCa-2 cells without BODIPY labeling. (B) CARS images from MIAPaCa-2 cells labeled with 4 μg/mL BODIPY for 1 hr. Scale bars: 20 µm.

**Figure S10.**
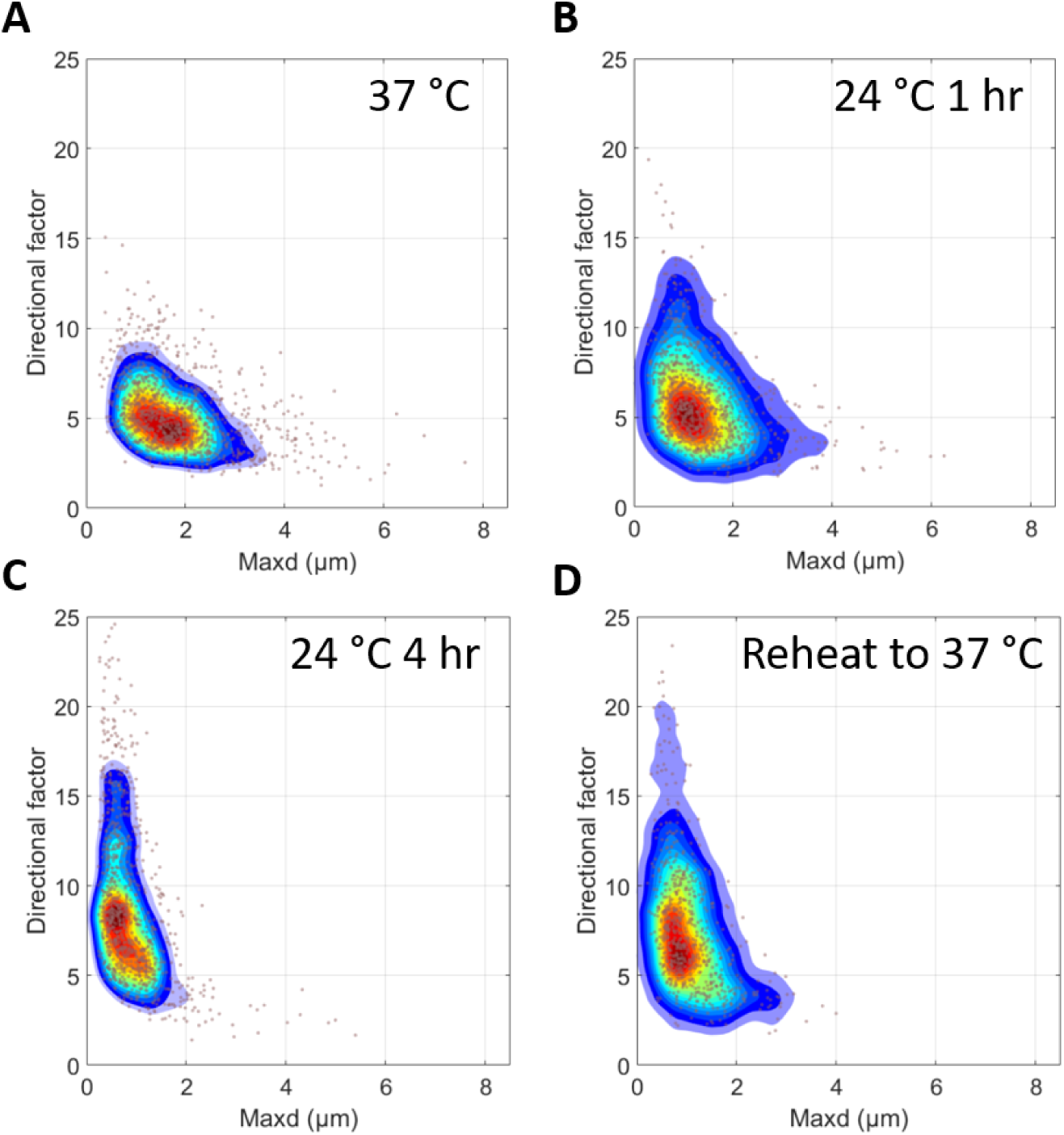
Long exposure to hypothermia caused irreversible changes in LD dynamics. (A)-(D) Dynamic signatures of MIAPaCa-2 cells at 37 °C, hypothermia exposure to 24 °C for 1 hr, hypothermia exposure to 24 °C for 4 hr, and reheat sample to 37 °C, projected onto the *DF-maxd* plane, respectively.

**Figure S11.**
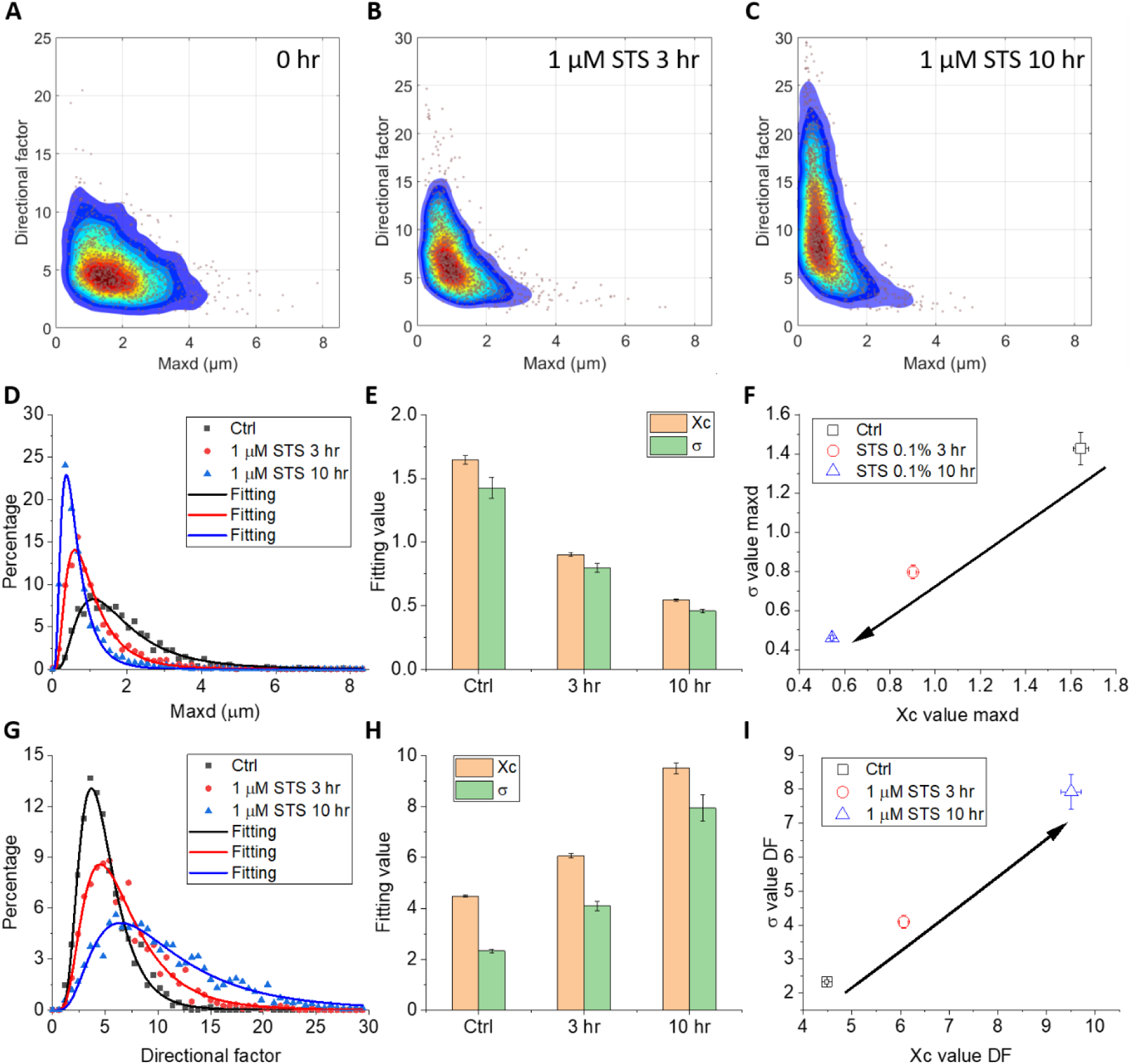
Changes of LD dynamics during apoptosis induced by 1 μM STS. (A)-(C) Dynamic signatures of MIAPaCa-2 control group and 1 μM STS treatment for 3 hr and 10 hr, respectively, projected onto the *DF-maxd* plane. (D) Histograms (dots) and the lognormal fitting of the histograms (solid curves) of MIAPaCa-2 cells at *maxd* domain from the control (black), 3 hr STS treatment (red), and 10 hr STS treatment (blue). (E) *Xc* and *σ* values derived from the lognormal fitting from each condition in panel (D). (F) Time-dependent STS treatment results plotted in the *maxd Xc-σ* domain. The arrow indicates the change over STS treatment time. (G) Histograms (dots) and the lognormal fitting of the histograms (solid curves) of MIAPaCa-2 cells at *DF* domain from the control (black), 3 hr STS treatment (red), and 10 hr STS treatment (blue). (H) *Xc* and *σ* values derived from the lognormal fitting from each condition in panel (G). (I) Time-dependent STS treatment results plotted in the *DF Xc-σ* domain. The arrow indicates the change over STS treatment time.

**Figure S12.**
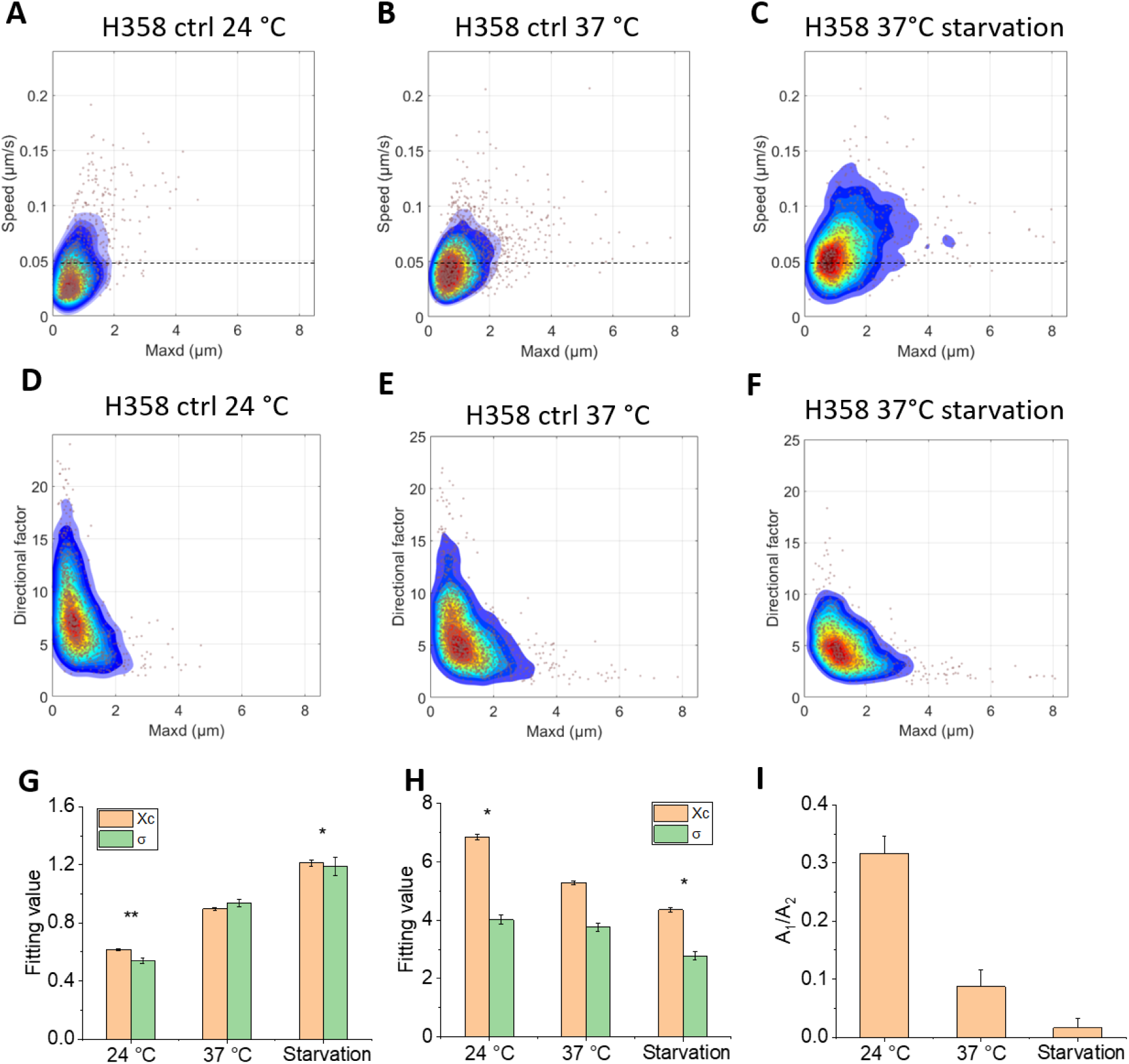
The dynamic signatures of H358 cells under hypothermia exposure and starvation conditions. (A)-(C) Dynamic signatures of H358 cells projected onto the *speed-maxd* plane during 24 °C hypothermia exposure for 1 hr, at normal culture condition, and after 24 hr starvation without glucose and FBS in the culture medium, respectively. (D)-(F) Dynamic signatures of H358 cells in the same three conditions as in (A)-(C), projected onto the *DF-maxd* plane. (G) *Xc* and *σ* values derived from the lognormal fitting of *maxd* from each condition in panels (A)-(C). (H) *Xc* and *σ* values derived from the lognormal fitting of *DF* from each condition in panels (D)-(F). (I) The ratio of amplitudes A_1_/A_2_ derived from the dual-lognormal fitting of the *speed* histograms from the three conditions.

**Figure S13.**
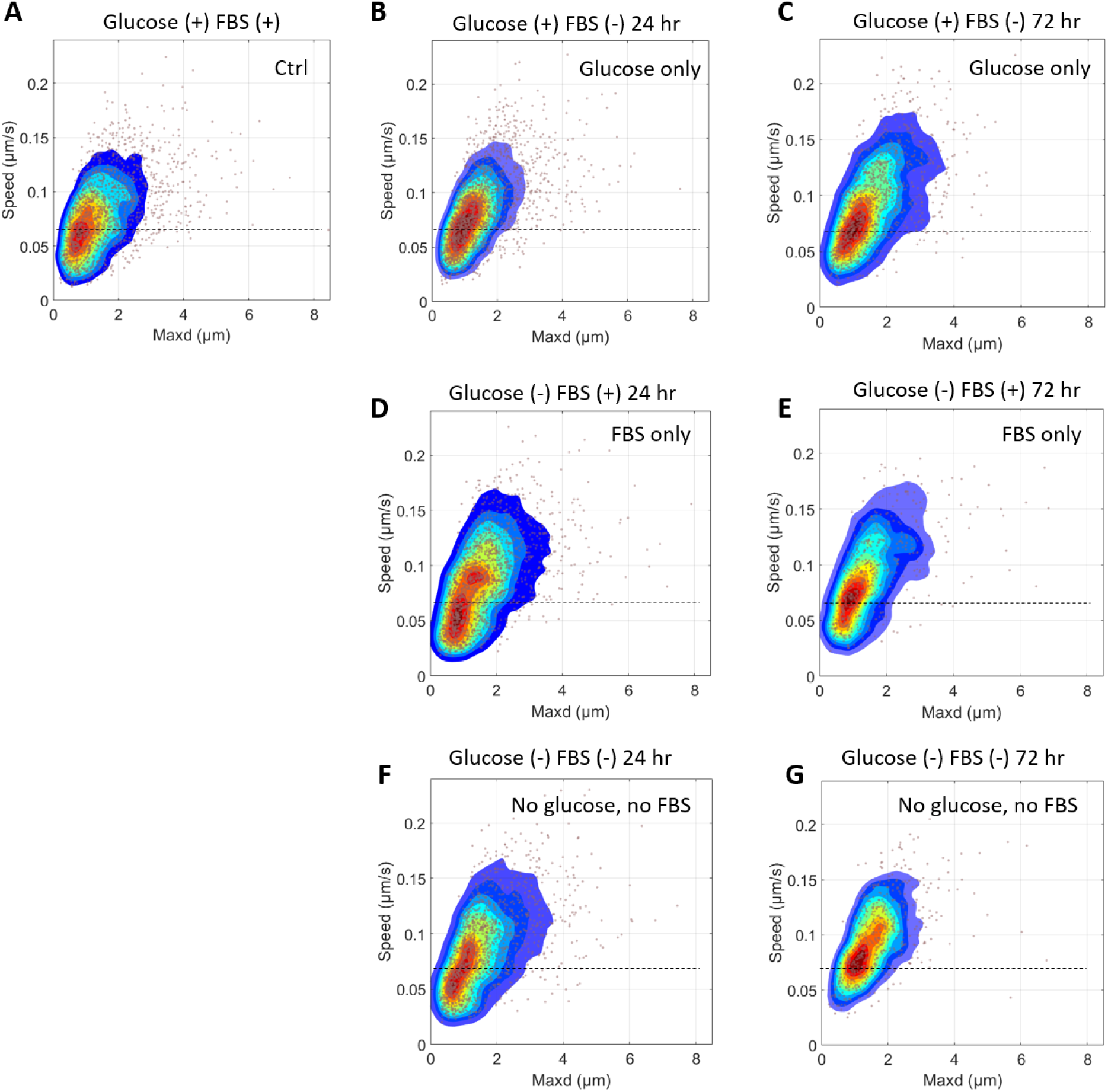
LD dynamic signatures in different starvation conditions. Dynamic signatures of MIAPaCa-2 cells projected onto the *speed-maxd* plane for (A) the control group; (B) and (C) with only glucose for 24 and 72 hr, respectively; (D) and (E) with only FBS for 24 and 72 hr, respectively; (F) and (G) with no glucose and no FBS for 24 and 72 hr, respectively. The dotted lines are shown as reference marks.

**Figure S14.**
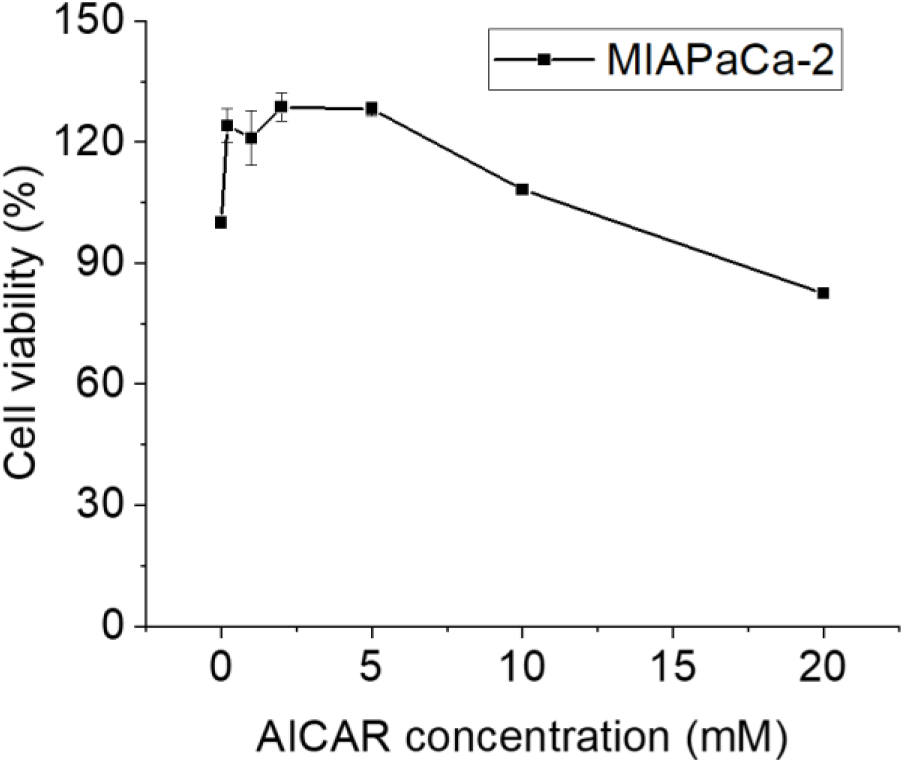
MTT viability assay of MIAPaCa-2 cells treated by AICAR (24 hr). MTT viability results for MIAPaCa-2 cells treated with various concentrations of AICAR in normal culture media for 24 hr. The viability was normalized with the control group (0 mM AICAR). n=6.

**Figure S15.**
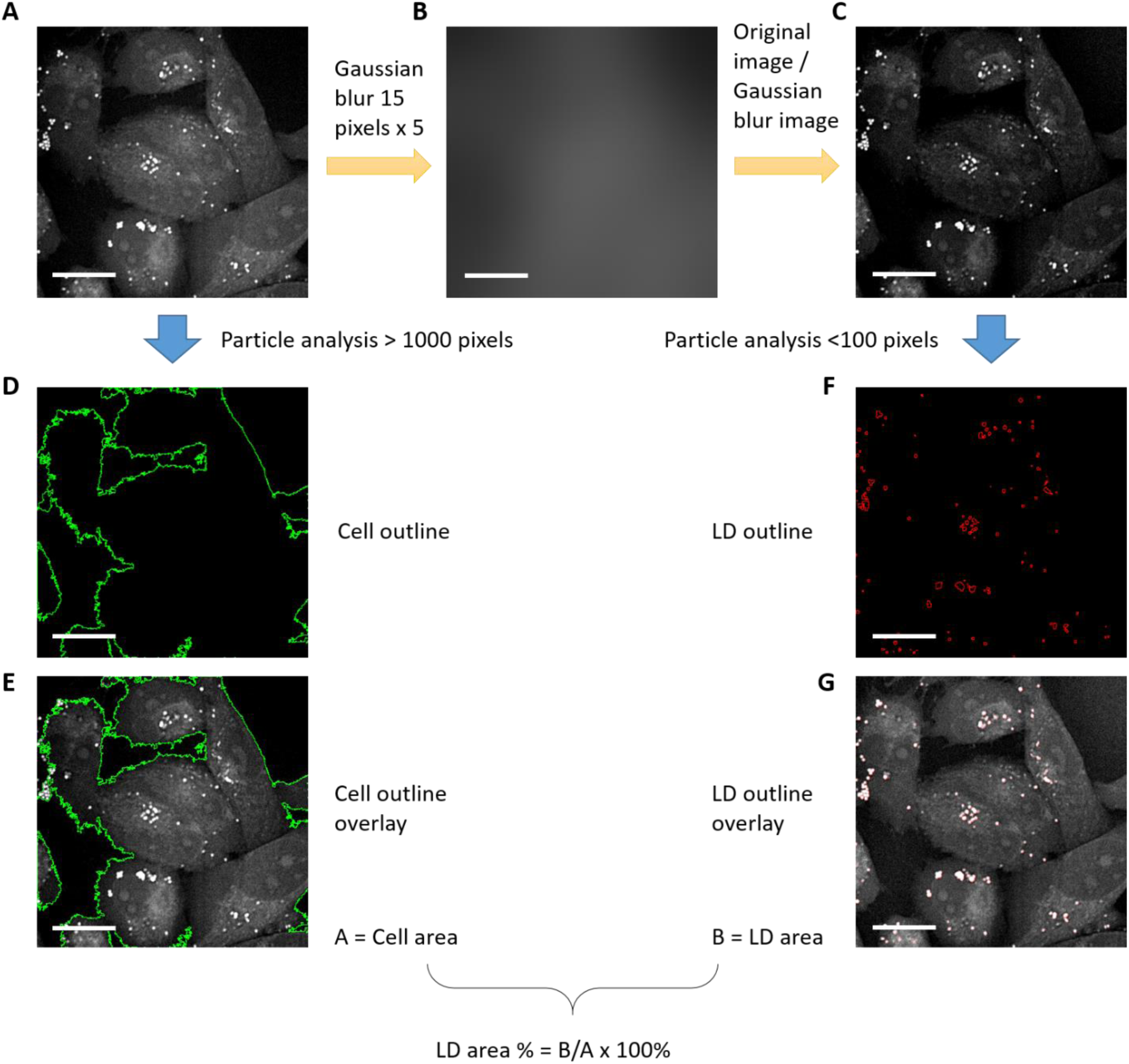
Quantitative analysis of LD area percentage. (A) The original CARS image. (B) Gaussian-blurred image using 15 pixels for 5 times. (C) Image obtained by dividing the original CARS image by the Gaussian-blurred image. (D) Particle tracking after intensity thresholding of panel (A) using particle size > 1000 pixels delineates the outlines of cells. (E) Superposition of image in panel (A) and the outlines in panel (D). (F) Particle tracking after intensity thresholding of panel (C) using particle size < 100 pixels outlines the LD particles. (G) Superposition of image in panel (A) and the outlines in panel (F). The total areas A for cells and B for LDs are calculated. The LD area % is defined as B/A x 100%. Scale bars: 20 µm.

**Figure S16.**
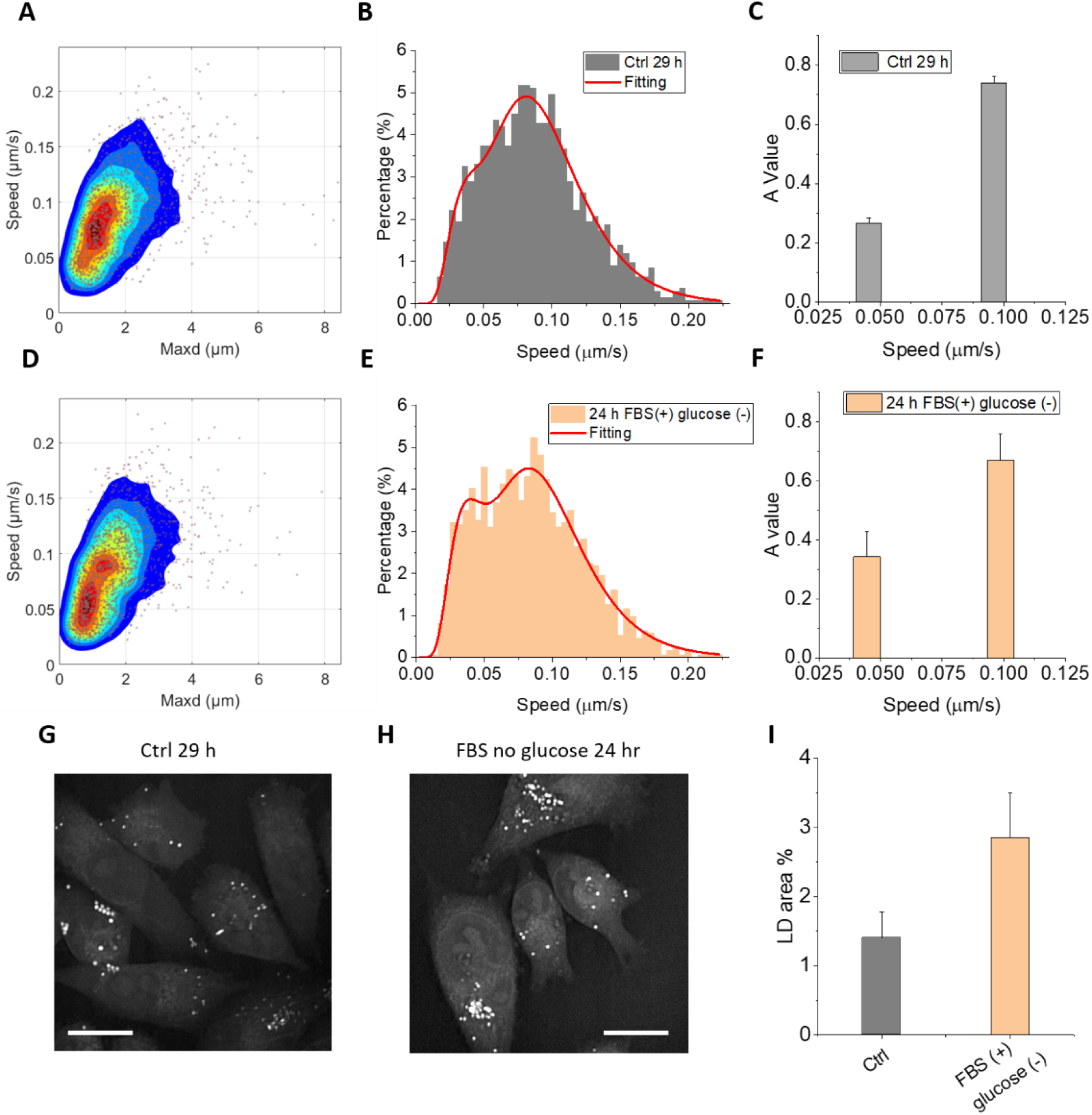
The changes of dynamic signatures and LDs of MIAPaCa-2 cells with only FBS as the energy source. (A) The dynamic signature of MIAPaCa-2 cells projected onto the *speed-maxd* plane for the control group. (B) The histogram (bars) in the *speed* domain and the dual-lognormal function fitting (solid curve) for the control group. (C) Amplitude values as a function of *Xc* derived from the histogram fitting in panel (B). (D) The dynamic signature of MIAPaCa-2 cells projected onto the *speed-maxd* plane after 24 hr culture with FBS and glucose-free DMEM. (E) The histogram (bars) in the *speed* domain and the dual-lognormal function fitting (solid curve) for the FBS-only cultured group. (F) Amplitude values as a function of *Xc* derived from the histogram fitting in panel (E). (G) and (H), sample CARS images for the control and FBS-only groups, respectively. (I) LD area % derived from the CARS images using the method described in Figure S15. Scale bars: 20 µm.

**Figure S17.**
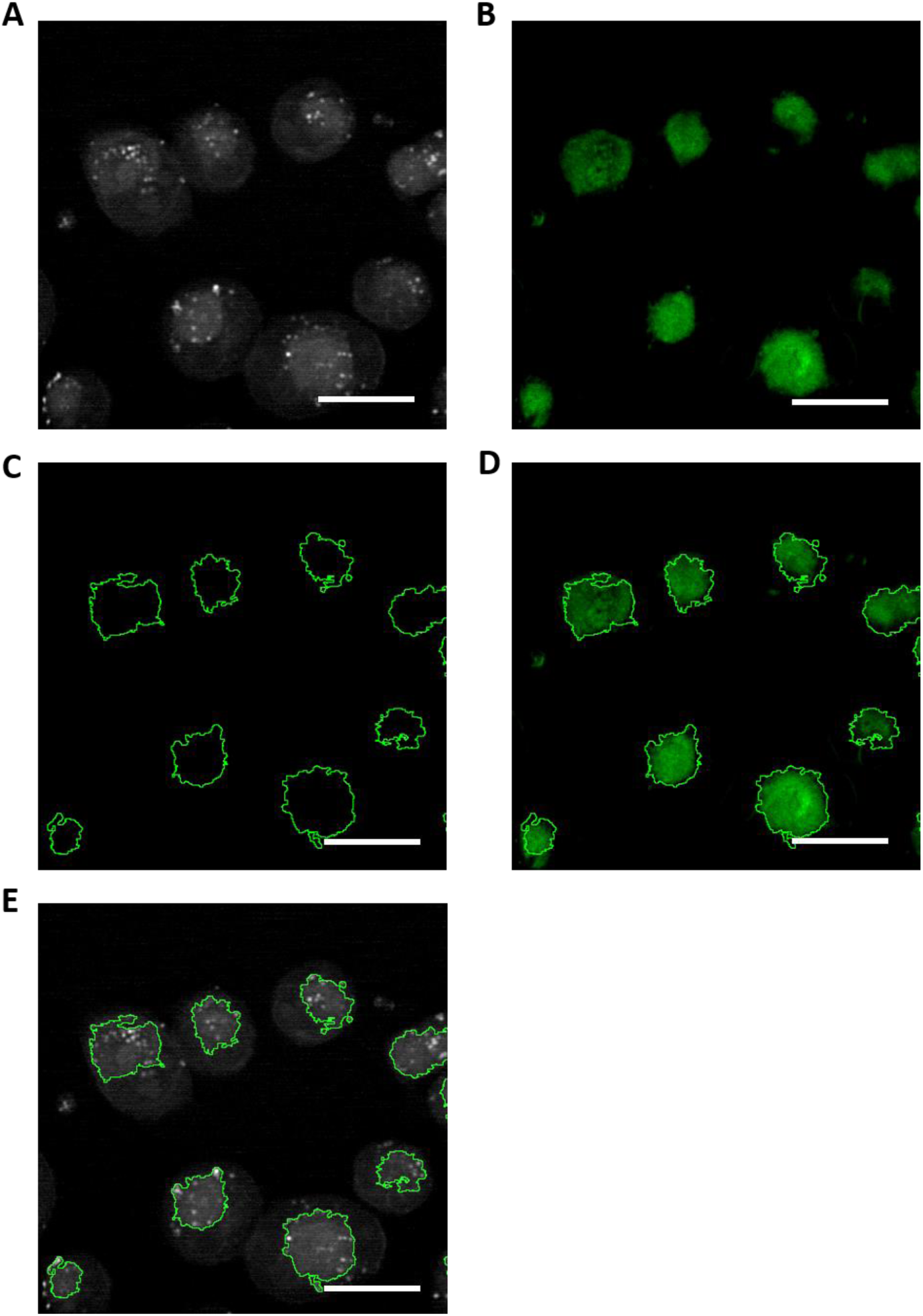
Correlate CARS intensity thresholding with ER Tracker™ labeling. (A) The original CARS image of MIAPaCa-2 cells. (B) TPEF image of ER from the same field of view in panel (A). The cells were labeled by ER Tracker™ Green. (C) Intensity thresholding from the original CARS image outlines ER areas. (D) Superposition of label-free CARS-outlined ER area and ER Tracker™ -labeled ER areas. (E) Superposition of the original CARS image with the ER outlines from panel (C).

**Figure S18.**
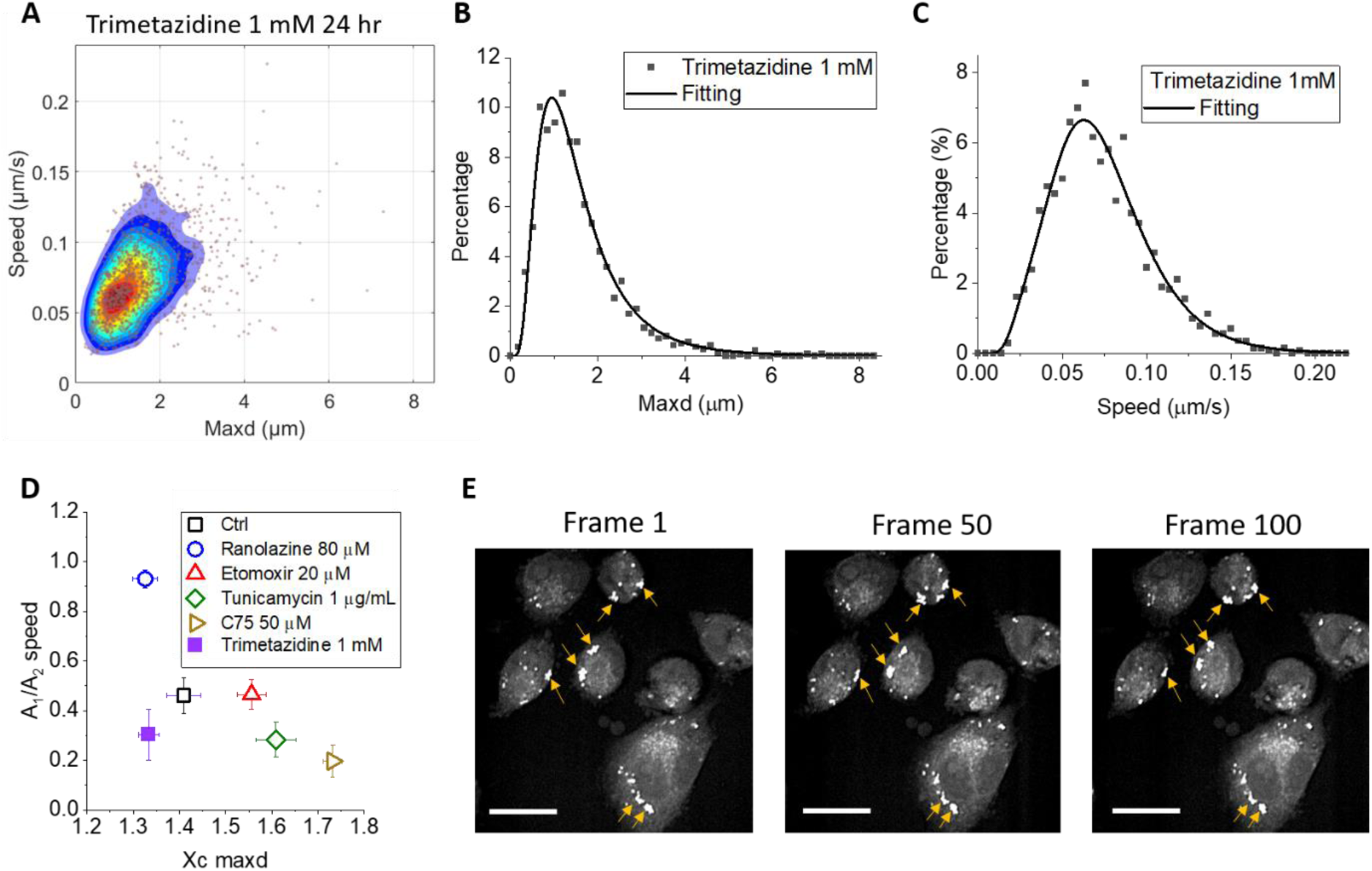
The impact of trimetazidine on LD dynamics. (A) The dynamic signature of MIAPaCa-2 projected onto the *speed-maxd* plane after 1 mM trimetazidine treatment for 24 hr. (B) The histogram (dots) in the *maxd* domain and the lognormal fitting (solid curve) for the trimetazidine treatment. (C) The histogram (dots) in the *speed* domain and the dual-lognormal function fitting (solid curve) for the trimetazidine treatment. (D) The dynamic feature of the trimetazidine treatment, together with other treatments, in the A_1_/A_2_ – *Xc* domain. (E) CARS images for the trimetazidine-treated MIAPaCa-2 cells. The time interval between 1-50 and 50-100 frames are 110 s. Arrows point out LD aggregations in each frame. Scale bars: 20 µm.

### Supplementary Videos

**Video S1.**
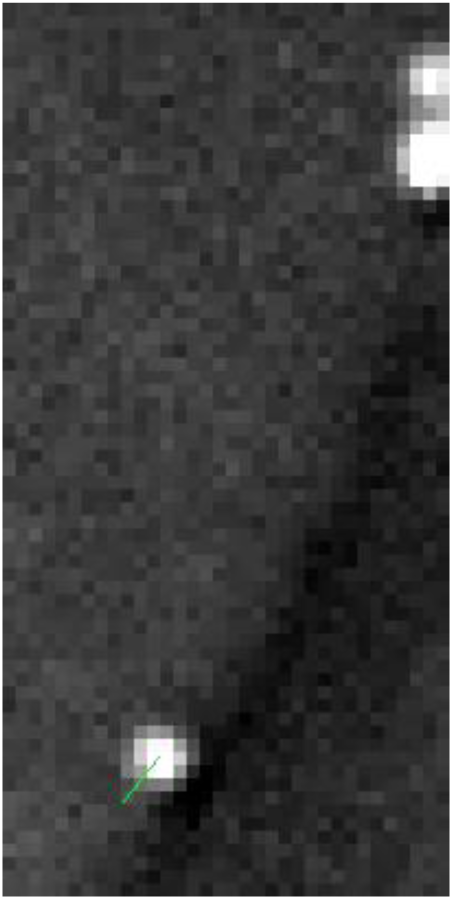
Tracing an LD trajectory in a living MIAPaCa2 cell shown in Figure 1C.

**Video S2.**
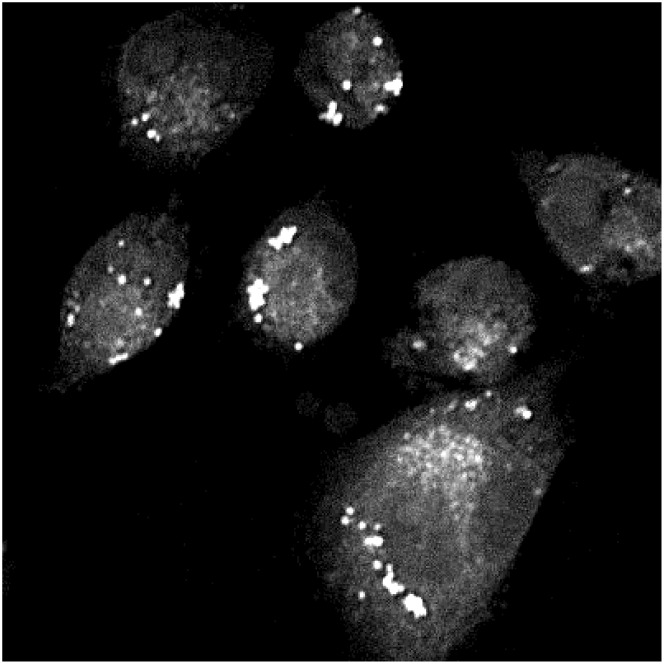
Time-lapse CARS images showing LD aggregations in MIAPaCa2 cells treated with 1 mM trimetazidine.

